# *Leishmania* infection triggers hepcidin-mediated proteasomal degradation of Nramp1 resulting in increased phagolysosomal iron availability

**DOI:** 10.1101/820340

**Authors:** Sourav Banerjee, Rupak Datta

## Abstract

Natural resistance associated macrophage protein 1 (Nramp1) was discovered as a genetic determinant of resistance against multiple intracellular pathogens, including *Leishmania*. It encodes a transmembrane protein of the phago-endosomal vesicles, where it functions as an iron transporter. But how Nramp1 expression is regulated in an infected macrophage is unknown. Its role in controlling iron availability to the intracellular pathogens and in determining the final outcome of an infection also remains to be fully deciphered. Here we report that Nramp1 protein abundance undergoes temporal changes in *Leishmania major* infected macrophages. At 12 hours post infection, there was drastic lowering of Nramp1 level accompanied by increased phagolysosomal iron availability and enhanced parasite growth. *Leishmania* infection-induced downregulation of Nramp1 was found to be caused by ubiquitin-proteasome degradation pathway. In fact, blocking of Nramp1 degradation with proteasome inhibitor resulted in depletion of phagolysosomal iron pool with significant reduction in the number of intracellular parasites. Further, we uncovered that this degradation process is mediated by the iron regulatory peptide hormone hepcidin that binds to Nramp1. Interestingly, Nramp1 protein level was restored to normalcy after 30 hours of infection with a concomitant drop in the phagolysosomal iron level, which is suggestive of a host counter defense strategy to deprive the pathogen of this essential micronutrient. Taken together, our study implicates Nramp1 as a central player in the host-pathogen battle for iron. It also unravels Nramp1 as a novel partner for hepcidin. The hitherto unidentified ‘hepcidin-Nramp1 axis’ may have a broader role in regulating macrophage iron homeostasis.

**Importance:** *Leishmania* parasites are the causative agents of a group of neglected tropical diseases called leishmaniasis. They reside within the phagolysosomes of mammalian macrophages. Since iron is an essential micronutrient for survival and virulence, intracellular *Leishmania* must acquire it from the tightly regulated macrophage iron pool. How this challenging task is accomplished remains a fundamental question in *Leishmania* biology. We report here that *Leishmania major* infection caused ubiquitin-proteasome-mediated degradation of natural resistance associated macrophage protein 1 (Nramp1). Nramp1 being an iron exporter at the phago-endosomal membrane, its degradation resulted in increased phagolysosomal iron availability thereby stimulating parasite growth. We also uncovered that Nramp1 degradation is controlled by the iron regulatory peptide hormone hepcidin. Interestingly, at a later stage of infection, Nramp1 protein level was restored to normalcy with simultaneous depletion of phagolysosomal iron. Collectively, our study implicates Nramp1 as a central player in the host-pathogen struggle for acquiring iron.

## Introduction

*Leishmania* belongs to the trypanosomatid group of protozoan parasites and they are the causative agents for a spectrum of human diseases collectively known as leishmaniasis. Depending on the *Leishmania* species, severity of the disease varies from self-resolving skin ulcers to life threatening infection of the visceral organs. With about 12 million currently affected people from nearly 100 endemic countries and more than 1 million new cases every year, leishmaniasis imposes a significant challenge to the global healthcare system (1, 2). Unavailability of a vaccine, limited chemotherapeutic options and increasing signs of drug resistance further aggravated the problem (3). Better understanding of *Leishmania* physiology and the mechanisms which enable the parasite to survive in host environment is therefore required to conceive novel strategies for effectively combating this disease.

*Leishmania* promastigotes are transmitted to humans by bite of a sand fly. Thereafter, they are rapidly phagocytosed by the macrophages either directly or via engulfment of parasite containing apoptotic neutrophils (4). Upon internalization, the parasites are delivered to the phagolysosome where they differentiate into the amastigote form and continue to proliferate till the cell bursts (5). While traversing through the phagosome maturation pathway, *Leishmania* has to overcome multitude of host-induced stress factors, including oxidative and nitrosative stresses, before encountering the low pH and hydrolytic enzyme-rich environment of the phagolysosome. Apart from facing these challenges, *Leishmania* amastigotes also have to scavenge all the essential nutrients from resource-limited phagolysosomal environment (6, 7). To subvert such adversaries, *Leishmania* parasites encode various nutrient acquisition genes and are also armored with defenses against free radicals, intracellular acidosis and lysosomal hydrolases (5, 8–11). Moreover, intracellular *Leishmania* also have the distinctive ability to manipulate host gene expression to adapt itself to the harsh environment (12–15). However, there has been very little effort to understand the molecular basis of these reprogramming events in *Leishmania* infected macrophages and to exploit them for anti-leishmanial drug discovery.

Genetic makeup of the host is also a critical factor in determining susceptibility to leishmaniasis and severity of the symptoms (16, 17). Linkage analysis and genome wide association studies led to identification of many disease modifier genes or genetic loci in mouse as well as in human (18, 19). Most prominent among them is the natural resistance associated macrophage protein 1 (Nramp1), which was originally identified as the *Bcg*/*Ity*/*Lsh* locus in the mouse chromosome 1. Mice expressing Nramp1 were shown to be resistant to diverse group of pathogens, such as *Mycobacteria*, *Salmonella* and *Leishmania* (20). Furthermore, data from human population-based studies demonstrated association between Nramp1 polymorphism and susceptibility to wide varieties of infectious diseases, including visceral and cutaneous leishmaniasis (21–23). Nramp1 belongs to the solute carrier protein family 11 (hence also known as SLC11A1) and is predicted to be an integral membrane protein with 12 transmembrane helices (24). It is exclusively expressed in the phagocytes where it was found to be localized in the lysosomes/late endosomes as well as in the membrane of maturing phagosomes/phagolysosomes (25, 26). Interestingly, a naturally occurring point mutation (G169D) in the fourth transmembrane domain of Nramp1 prevented proper maturation of the protein and the mice harboring this mutation were vulnerable to different types of infections (20, 27). Functional characterization of Nramp1 uncovered its role as an iron transporter at the phagosomal membrane, however, the direction of iron trafficking remained a matter of controversy (24, 28). Increased cytoplasmic influx of iron observed in Nramp1 expressing cells supported the role of Nramp1 in mobilizing iron from phagosomal compartment into the cytoplasm. This led to the hypothesis that functional Nramp1 restricts pathogen growth by depriving them of iron, which is an essential micronutrient (29, 30). However, there are contradictory reports suggesting Nramp1 imports iron into the phagosomes where it catalyzes generation of hydroxyl radical and thereby inflicting oxidative damage to the pathogens (31, 32). Thus, despite its proven role in conferring resistance against a wide variety of pathogenic infections, there is still some ambiguity regarding the mode of action of Nramp1. Also, whether Nramp1 status is modulated during the course of an infection and how its function influences the outcome of host pathogen interaction remains completely unknown. In this work we addressed these unresolved issues in a macrophage infection model for *Leishmania major*. We report here that *Leishmania* infection results in ubiquitin-proteasome-mediated degradation of macrophage Nramp1 at 12 hours post infection followed by almost complete recovery of Nramp1 level after 30 hours. Nramp1 downregulation at 12 hours post infection led to increased availability of iron within the phagolysosomes and enhanced intracellular parasite growth. Interestingly, our work also revealed that hepcidin, an iron regulatory peptide hormone, binds with Nramp1 and facilitates its degradation (33). Thus, besides identifying a hitherto unknown strategy by which intracellular *Leishmania* modulates host’s iron withholding machinery for its own survival benefit, this study unraveled the role of hepcidin-Nramp1 signaling axis in regulating the macrophage iron recycling process.

## Results

### Nramp1 is recruited to the phagolysosomes in *L. major* infected macrophages

Although Nramp1 is known to mediate resistance against various intracellular pathogens, whether its own status is altered during the course of infection is unknown. To address this, we first sought to compare subcellular distribution of Nramp1 in uninfected versus *L. major* infected macrophages by immunofluorescence staining. It was revealed that in uninfected J774A.1 cells (a BALB/c mouse derived macrophage cell line), Nramp1 was distributed in punctate intracellular structures, which colocalized strongly with Rab11 positive endocytic vesicles (PCC: 0.93±0.028) and to a lesser extent with the lysosomal marker, Lamp1 (PCC: 0.43±0.06). Whereas, in *L. major* infected macrophages, Nramp1 was found to reside mostly in the Rab11/Lamp1 double-positive vesicles (PCC for Rab11: 0.89±0.05, PCC for Lamp1: 0.67±0.09) at 12 hours post infection (Fig. 1A to D). These results suggest that Nramp1 is preferentially recruited to the phagolysosomal compartment upon *L. major* infection. It is also worth noting that there was a marked reduction in the Nramp1 protein level in *L. major* infected macrophages as compared to their uninfected counterparts (Fig. 1A and B). This intriguing finding prompted us to undertake a systematic analysis of the relative abundance of Nramp1 during the course of *L. major* infection.

**FIG 1.**
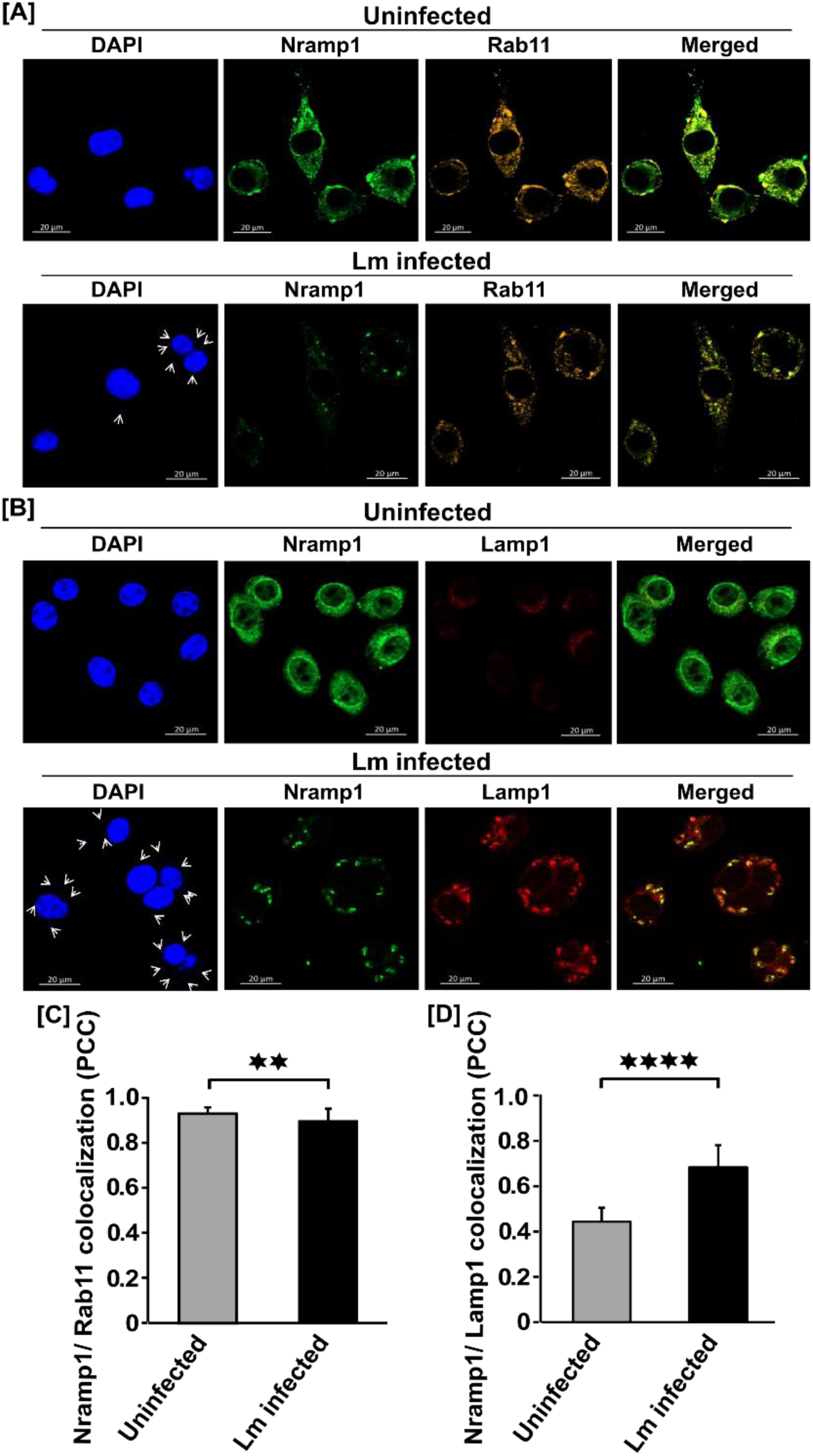
Nramp1 is recruited to the phagolysosomes in *L. major* infected macrophages. (A and B) J774A.1 macrophages were infected with *L. major* (Lm) and co-immunostained with anti-Nramp1 (green) and anti-Rab11 (cyan red) (A) or anti-Nramp1 (green) and anti-Lamp1 (red) (B) at 12hrs post infection. The merged images represent colocalization of the proteins (yellow). Nuclei were stained with DAPI (blue). Uninfected macrophage cells were used as control. White arrows indicate the presence of intracellular parasites in infected cells (smaller nuclei). Cells were visualized with Zeiss Apotome microscope using 63X oil immersion objective. Scale bar: 20µm. (C and D) Bar diagrams represents Pearson’s colocalization coefficient (PCC) of Nramp1 with Rab11 (C) or with Lamp1 (D) measured using ImageJ software. Grey bar represents uninfected macrophages whereas black bar represents Lm infected macrophages. At least 20 cells from three independent experiments were scored in each of the cases. Error bars represent standard error mean (SEM). **, p≤0.01; ****, p≤0.0001.

### Temporal modulation of Nramp1 protein level in *L. major* infected macrophages

Following the lead from the previous experiment, we infected J774A.1 macrophages with *L. major* promastigotes and then Nramp1 protein level was visualized by immunofluorescence staining at different time points after infection (2 – 30 hours). Although Nramp1 expression remained almost unaltered at early time points after infection (2 or 6 hours), there was a significant reduction in its level (by > 50%) at 12 hours post infection. Interestingly, the level of Nramp1 was almost restored to normalcy at a later time point i.e. 30 hours post infection (Fig. 2A to D). The status of Nramp1 in J774A.1 macrophages during the course of infection was independently verified by western blot analysis, the results of which are in complete agreement with our immunofluorescence data (Fig. S1). Before proceeding further, we also wanted to validate our cell line-based data in a primary macrophage cell. For this we isolated thioglycolate elicited peritoneal macrophages from BALB/c mice and infected those cells with *L. major* promastigotes. As seen earlier in J774A.1 cell line, Nramp1 protein level in *L. major* infected peritoneal macrophages underwent drastic reduction at 12 hours post infection but reverted almost to the normal level after 30 hours (Fig. S2). Collectively, these data provided unambiguous evidence that cellular abundance of the Nramp1 protein indeed changes during the course of infection, which is possibly an outcome of the complex dynamics of host pathogen interaction.

**FIG 2.**
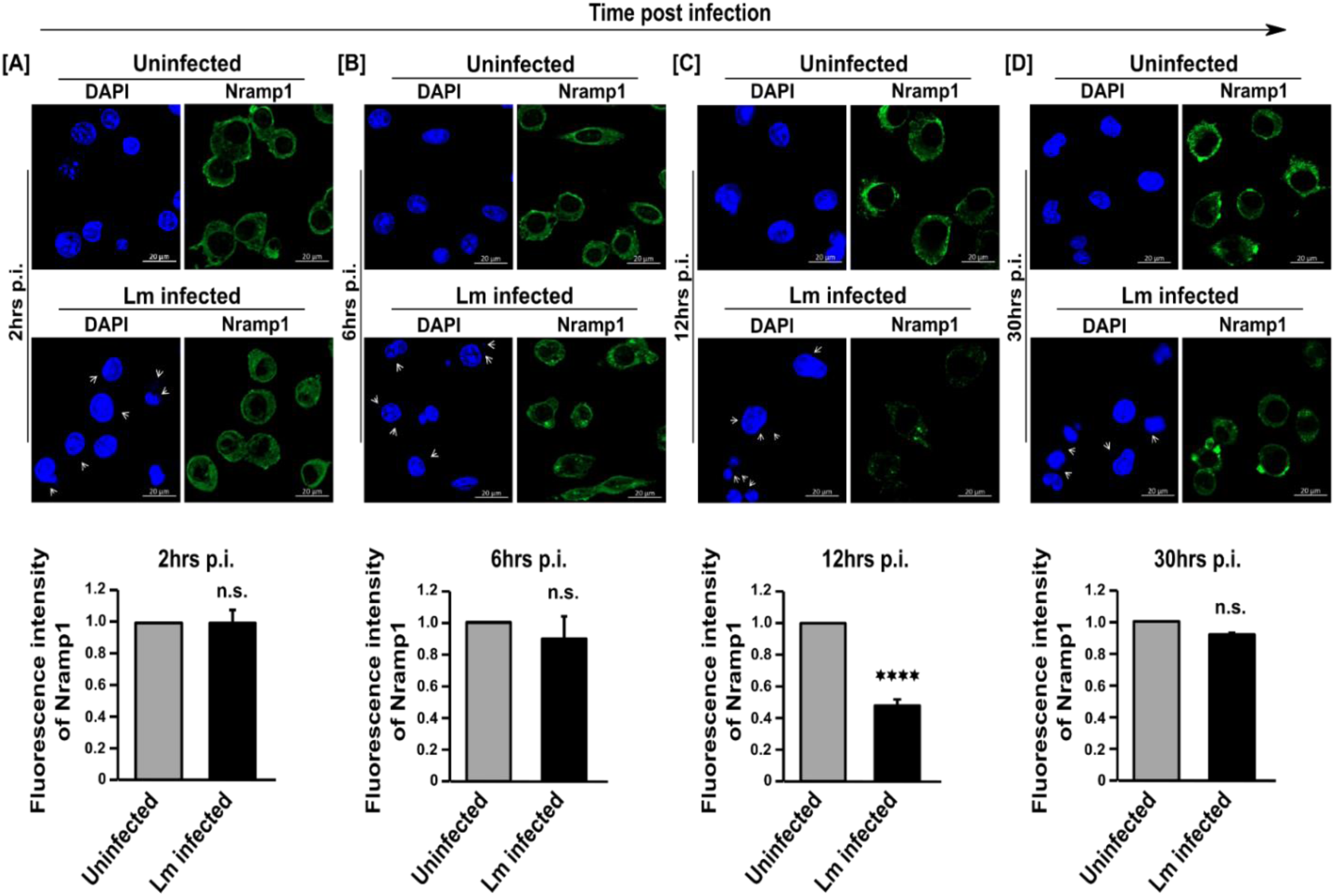
Alteration of Nramp1 protein level during the course of *L. major* infection. (A-D) Nramp1 protein level was visualized by immunostaining with anti-Nramp1 (green) in uninfected or *L. major* (Lm) infected J774A.1 macrophages at different time points post infection (2-30hrs). Nuclei were stained with DAPI (blue). Arrows in the Lm infected panel indicates the presence of intracellular parasites (smaller nuclei). Cells were visualized under 63X oil immersion objective of Zeiss Apotome microscope. Scale bar: 20µm. Lower panel represents the quantitative estimation of Nramp1 expression as indicated by the relative fluorescence intensity of Nramp1 at indicated time points in uninfected (grey bar) and Lm infected (black bar) macrophage cells. Several fields were imaged using Olympus IX-81 epifluorescence microscope and at least 100 macrophage cells were analyzed for the fluorescence intensity measurement using ZEN software of the microscope. Uninfected cells were used as reference sample during quantification. Error bars represent SEM calculated from three independent experiments. ****, p≤0.0001; n.s., non-significant.

### Downregulation of Nramp1 in *L. major* infected macrophages resulted in increased phagolysosomal iron content and higher intracellular parasite burden

Since Nramp1 is as a phagosomal/phagolysosomal iron transporter, the next obvious question was whether this significant drop in the Nramp1 level at 12 hours post infection has any functional implication in regulating iron availability within the phagosomal/phagolysosomal compartments (25). For this, we isolated Nramp1/Rab11 double positive phagosomal/ phagolysosomal fraction from both uninfected and *L. major* infected J774A.1 macrophages by sucrose density gradient centrifugation and measured iron content in them by ferrozine-based colorimetric assay (Fig. 3A to C). We observed that iron content in the phagosomes/phagolysosomes isolated from *L. major* infected macrophages at 12 hours post infection was nearly double than in those isolated from the uninfected cells (Fig. 3D). This correlation between decreased Nramp1 protein level and increased phagosomal/phagolysosomal iron concentration is consistent with the reported role of Nramp1 as a phagosomal iron exporter (29, 30). This was further validated when phagosomal/phagolysosomal iron content was measured at 30 hours post infection, the time point by which Nramp1 protein level was almost restored to normalcy. By this time there was a significant drop in the phagosomal/phagolysosomal iron concentration in the infected cells and it became almost comparable to that in the uninfected cells (Fig. 3D). Interestingly, at 12 hours post infection, increased phagosomal/phagolysosomal iron content coincided with maximum intracellular parasite burden measured over the time course of infection (Fig. 3E). This data reconfirmed the role of iron as a vital nutrient for *Leishmania* parasites residing in the phagolysosomal niche (34).

**FIG 3.**
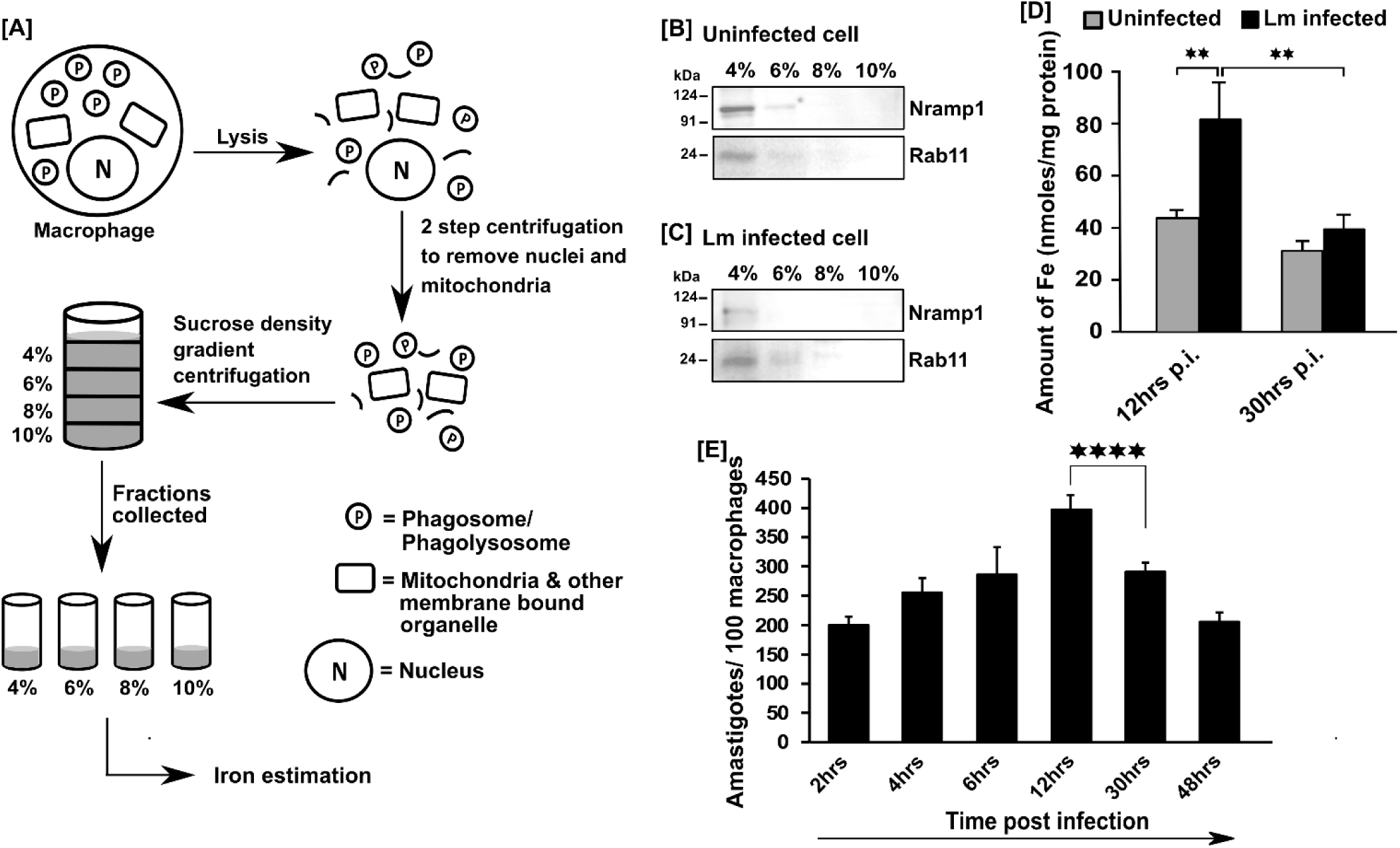
Downregulation of Nramp1 in *L. major* infected macrophages resulted in increased phagosomal/phagolysosomal iron content and higher intracellular parasite burden. (A) Schematic diagram showing subcellular fractionation to isolate phagosomes/phagolysosomes from macrophage cells using sucrose density gradient centrifugation followed by ferrozine-based iron estimation assay. (B and C) Western blot of the subcellular fractions prepared from uninfected (B) and *L. major* (Lm) infected (C) macrophage cells at 12hrs post infection (p.i.) using anti-Nramp1 and anti-Rab11 antibody to verify the presence of both Namp1 and Rab11 at their predicted molecular weight of ∼100 kDa and 24 kDa respectively. (D) Bar diagram representing phagosomal/phagolysosomal iron content as measured by ferrozine assay in uninfected (grey bar) and Lm infected (black bar) macrophage cells at 12hrs and 30hrs p.i. Error bars represents SEM values calculated from at least three independent experiments. (E) Intracellular parasite burden in Lm infected macrophage cells measured over 2-48hrs p.i. Intracellular parasite burden (amastigotes/ 100 macrophages) was quantified from at least 100 macrophage cells. Error bars represent SEM values calculated from at least three independent experiments. **, p≤0.01; ****, p≤0.0001.

### *L. major* infection caused Nramp1 degradation via ubiquitin-proteasomal pathway

To investigate the mechanism by which *L. major* infection causes downregulation of Nramp1, we first compared Nramp1 transcript levels in uninfected and *L. major* infected J774A.1 macrophages at 12 hours post infection. From our RT-qPCR data it is evident that transcription of Nramp1 remained unaltered, suggesting that its downregulation in *Leishmania* infected cells occurs via a post-transcriptional mechanism (Fig. 4A). We next examined the role of proteasomal activity in degradation of Nramp1. For this J774A.1 macrophages were treated with the proteasome inhibitor, MG132, immediately after infection with *L. major* and the Nramp1 protein level was visualized by immunofluorescence staining at 12 hours post infection (35). MG132 treatment completely prevented *Leishmania* infection-induced downregulation of Nramp1, implying that its degradation is mediated by the proteasomal machinery (Fig. 4B and C). This result was also independently validated by western blot analysis (Fig. S3). Since proteasomal substrates are usually ubiquitinated prior to degradation, we decided to check the ubiquitination status of Nramp1 in uninfected and *L. major* infected J774A.1 macrophages at 12 hours post infection (36). Whole cell lysate immunoprecipitation with anti-Nramp1 followed by western blot with ubiquitin antibody confirmed that ubiquitination of Nramp1 was significantly increased upon *Leishmania* infection thus making it an ideal target for proteasomal degradation (Fig. 4D).

**FIG 4.**
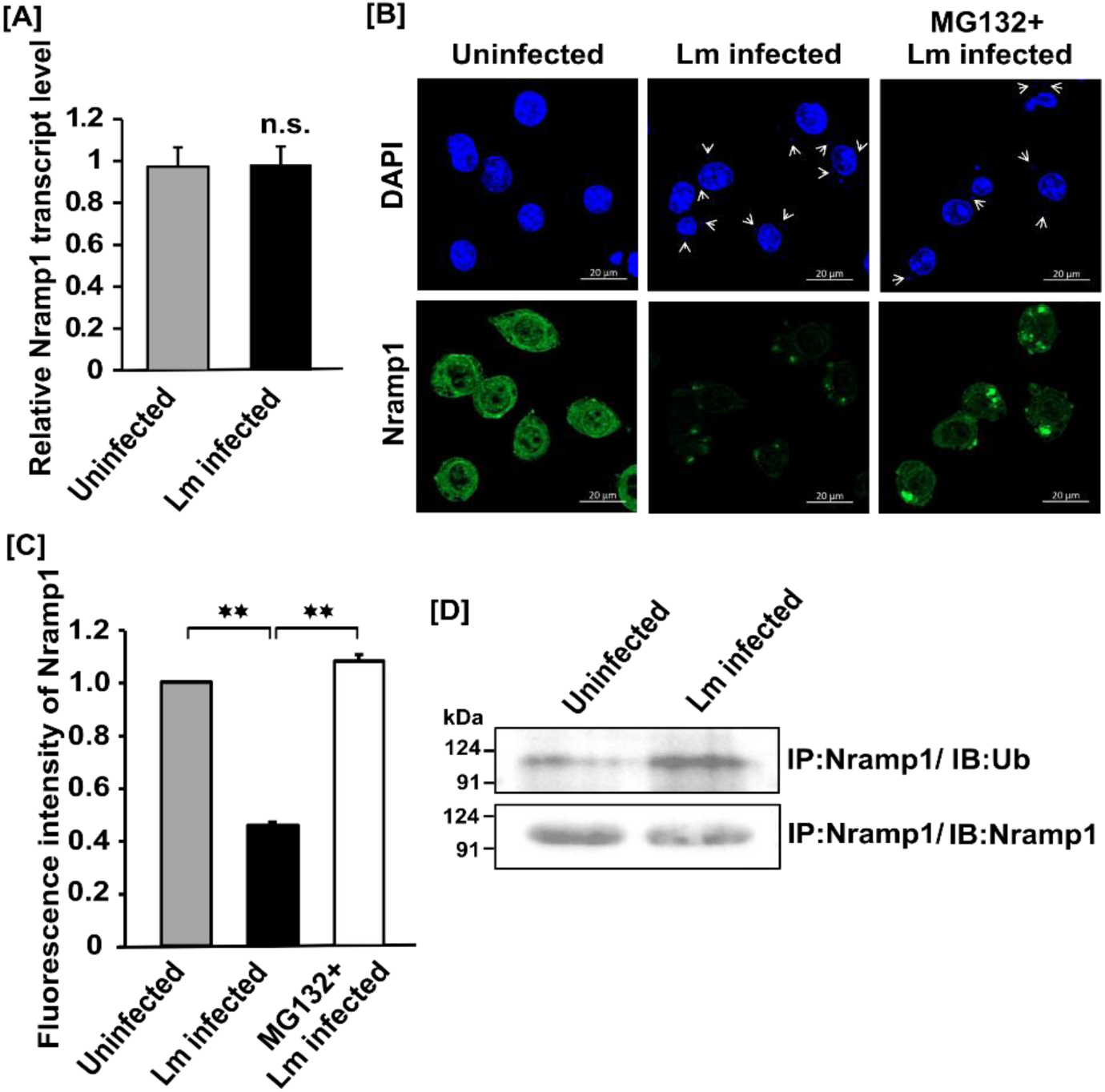
*L. major* infection causes Nramp1 degradation via ubiquitin-proteasomal pathway. (A) Representative bar diagram showing Nramp1 transcript level in uninfected (grey bar) and *L. major* (Lm) infected (black bar) macrophage cells at 12hrs post infection determined by RT-qPCR using β-Actin as endogenous control gene and uninfected cell as reference sample. Error bars represent SEM values calculated from 3 independent experiments. **, p≤0.01. (B) Nramp1 protein level was examined by immunostaining using anti-Nramp1 (green) in uninfected, Lm infected and 1µM MG132 treated Lm infected J774A.1 macrophage cells at 12hrs post infection. DAPI (blue) was used to stain *Leishmania* and macrophage cell nuclei. Arrows indicate the presence of intracellular parasites in Lm infected macrophages. Cells were visualized with Zeiss Apotome microscope using 63X oil immersion objective. Scale bar: 20µm. (C) Bar diagram depicts relative fluorescence intensity of Nramp1 in uninfected (grey bar), Lm infected (black bar) and 1µM MG132 treated Lm infected macrophage cells (white bar). Fluorescence intensity was measured from at least 100 macrophage cells imaged under Olympus IX-81 epifluorescence microscope using ZEN software. During quantification uninfected cells were considered as reference sample. Error bars represent SEM values calculated from three independent experiments. **, p≤0.01. (D) Western blot to verify the ubiquitination status of Nramp1 protein. Both uninfected and Lm infected J774A.1 macrophage cells at 12hrs post infection (p.i.) were lysed and cell lysates were analyzed for ubiquitination of Nramp1 by immuno-precipitation (IP) using anti-Nramp1 antibody followed by immunoblotting (IB) with anti-ubiquitin antibody (top panel). Uniform level of protein input was verified by IP as well as IB with anti-Nramp1 (lower panel).

### Nramp1 stabilization upon proteasomal inhibition resulted in decreased phagolysosomal iron content and lowering of intracellular parasite burden

Since the data presented in Fig. 3 indicated that Nramp1 exports iron from the phagosomes/phagolysosomes, we were prompted to check whether iron concentration in these compartments is decreased when Nramp1 in *L. major* infected macrophages is stabilized by inhibition of proteasomal activity. At 12 hours post infection, MG132-treated macrophages indeed had significantly lower phagosomal/phagolysosomal iron content than the infected cells that were not treated with the proteasomal inhibitor. In fact, phagosomal/phagolysosomal iron content in the MG132-treated, *L. major* infected macrophages was almost same as in the uninfected macrophages (Fig. 5A). Importantly, MG132 treatment also led to significant lowering of the intracellular parasite burden in the macrophages, which is likely to be a result of iron limiting phagolysosomal environment (Fig. 5B).

**FIG 5.**
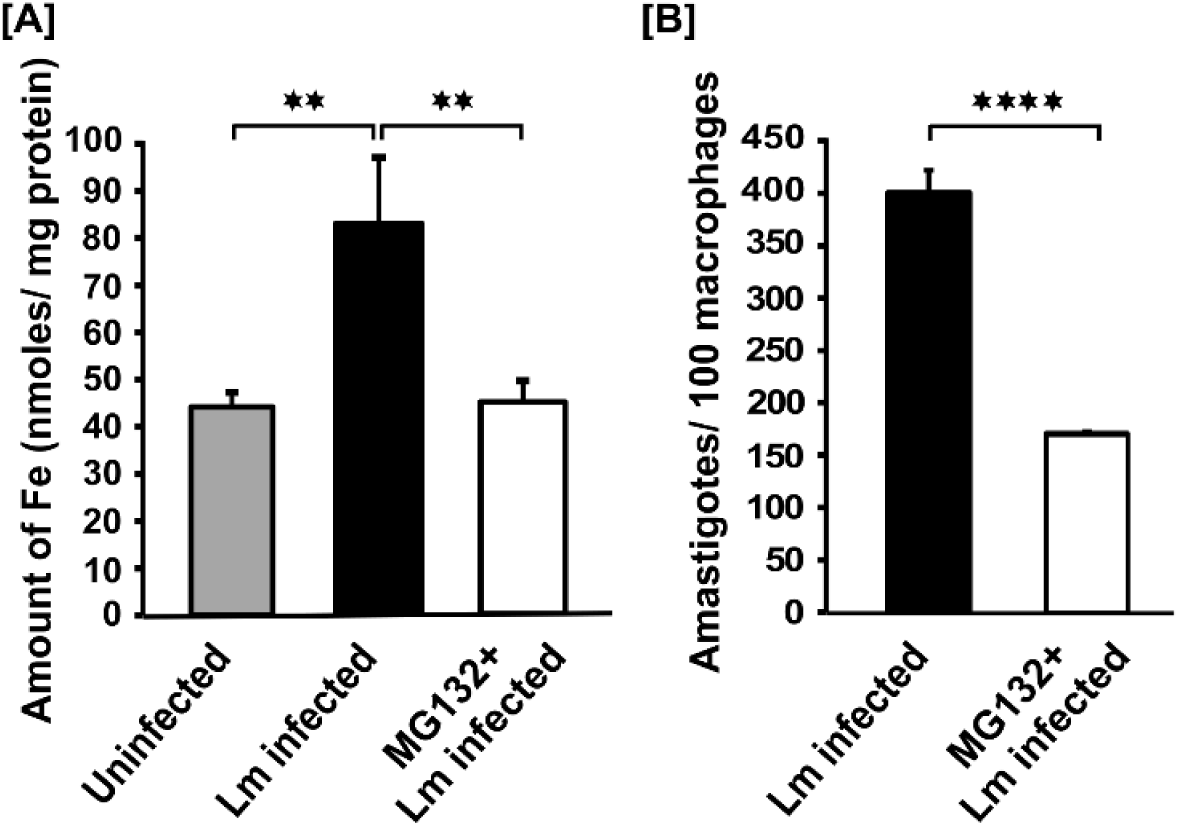
Nramp1 stabilization upon proteasomal inhibition resulted in decreased phagolysosomal iron content and lowering of intracellular parasite burden. (A) Bar diagram representing phagosomal iron content as measured by ferrozine assay in uninfected (grey bar), *L. major* (Lm) infected (black bar) and 1µM MG132 treated Lm infected (white bar) J774A.1 macrophage cells at 12hrs post infection as described earlier. Error bars represent SEM values calculated from at least three independent experiments. (B) Bar diagram showing amastigotes/ 100 macrophages count either in Lm infected (black) or in 1µM MG132 treated Lm infected (white) J774A.1 macrophage cells. Error bars represent SEM values calculated from at least 100 macrophage cells of three independent experiments. **, p≤0.01; ****, p≤0.0001.

### *L. major* infection-induced hepcidin surge in the macrophage is responsible for Nramp1 degradation, enrichment of phagolysosomal iron pool and enhanced parasite growth

Infection of macrophages with *L. amazonensis* was earlier shown to induce transcription of the iron regulatory peptide hormone hepcidin, which in turn caused degradation of the cell surface iron exporter ferroportin. This led to increased intracellular parasite growth, presumably due to enhanced macrophage iron content (37). However, ferroportin downregulation would increase the cytosolic iron pool, which cannot be directly accessed by the *Leishmania* parasites residing within the phagolysosomal compartment (37–39). So, it remained unclear how *Leishmania* infection-induced upregulation of hepcidin could stimulate intracellular parasite growth. Since Nramp1 closely resembles ferroportin in terms of its iron exporting property and membrane topology with 12 transmembrane helices, we decided to check if its expression is also regulated by hepcidin (24, 40). For this, we first validated the status of hepcidin expression in *L. major* infected J774A.1 macrophages. Consistent with the earlier results obtained with *L. amazonensis* infection, we observed sharp increase in hepcidin mRNA and protein levels in the *L. major* infected macrophages as compared to their uninfected counterparts (Fig. S4A and B) (37). It is worth noting that in the infected macrophages, hepcidin was found to be mostly distributed in vesicular compartments suggesting that it may have important intracellular role(s) in addition to its well-known paracrine action (Fig. S4B) (41). To check if Nramp1 expression is indeed regulated by intracellular hepcidin, we decided to treat the cells with heparin, an established transcriptional blocker for hepcidin that restricted hepcidin up-regulation in *L. major* infected macrophages (Fig. 6A) (42). Interestingly, treatment of the macrophages with heparin provided complete protection against *L. major* infection-induced downregulation of Nramp1 protein level without affecting cell viability as revealed by our immunofluorescence data as well as western blot analysis (Fig. 6B and C; and Fig. S5A to C). Increased Nramp1 level in heparin treated macrophages was accompanied by significantly reduced phagosomal iron content, which resulted in drastic lowering of intracellular parasite burden (Fig. 6D and E). Co-immunoprecipitation assay further demonstrated physical interaction of hepcidin with Nramp1 indicating that hepcidin binding may induce ubiquitination and proteasome-mediated degradation of Nramp1 similar to what has been earlier reported for ferroportin (Fig. 6F) (43, 44). Taken together our data strongly suggest that Nramp1 degradation in *L. major* infected macrophages occurs via hepcidin-dependent autocrine pathway, which is critical for maintaining an iron rich phagolysosomal environment required for parasite growth.

**FIG 6.**
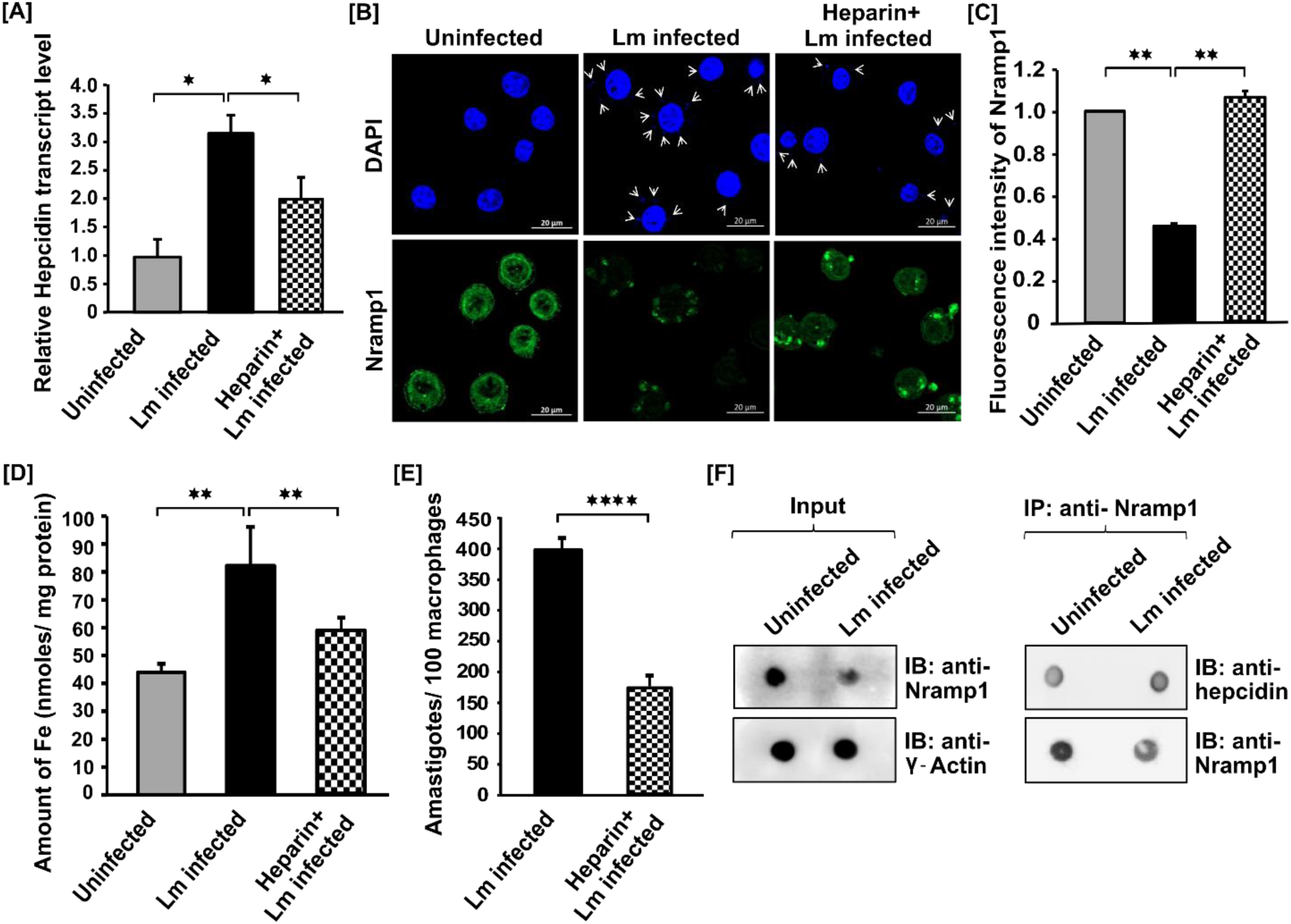
*L. major* infection-induced hepcidin upregulation in macrophage is responsible for Nramp1 degradation. (A) Bar diagram showing RT-qPCR data of hepcidin transcript level in uninfected (grey bar), *L. major* (Lm) infected (black bar) and 4µg/ml heparin treated Lm infected J774A.1 macrophage cells (stripped bar) at 12hrs post infection. All the measurements were performed using uninfected cell as reference sample and β-actin as an endogenous control gene. Error bars represent SEM values calculated from three independent experiments. *, p≤0.05. (B) Nramp1 protein level was observed by immunostaining using anti-Nramp1 (green) in uninfected, Lm infected and 4µg/ml heparin treated Lm infected J774A.1 macrophage cells at 12hrs post infection. *Leishmania* and macrophage nuclei were stained with DAPI (blue). Arrows indicate the presence of intracellular parasites in Lm infected macrophages. Cells were visualized with Zeiss Apotome microscope using 63X oil immersion objective. Scale bar: 20µm. (C) Bar graph showing relative fluorescence intensity of Nramp1 measured using ZEN software from at least 100 macrophage cells of uninfected (grey bar), Lm infected (black bar) and heparin treated Lm infected macrophage cells (stripped bar) imaged under Olympus IX-81 epifluorescence microscope. Error bars represent SEM values calculated from three independent experiments. *, p≤0.05; **, p≤0.01. (D) Bar diagram showing phagosomal/phagolysosomal iron level as measured by ferrozine assay in uninfected (grey bar), *L. major* (Lm) infected (black bar) and 4µg/ml heparin treated Lm infected (stripped bar) J774A.1 macrophage cells at 12hrs post infection. (E) Intracellular parasite burden in Lm infected (black bar) and 4µg/ml heparin treated Lm infected (stripped bar) J774A.1 macrophage cells at 12 hrs p.i. Amastigotes/ 100 macrophages was measured from at least 100 macrophage cells of three independent experiments as detailed previously. Error bars represent SEM values calculated from the experiments. **, p≤0.01; ****, p≤0.0001. (F) Both uninfected and Lm infected J774A.1 macrophage cells at 12hrs post infection were lysed and cell lysates were immuno-precipitated (IP) using anti-Nramp1 antibody. Either whole cell lysates of those cells (input) in left panel or immunoprecipitated (IP) samples in the right panel were subjected to immunodot blot (IB) with anti-hepcidin and anti-Nramp1 to verify the presence of these proteins. Immunodot blot with anti-γ Actin served as loading control.

## Discussion

Influence of host genetic factors on the outcome of *Leishmania* infection is widely reported (16–19). Among all the infection modifier genes, Nramp1 is particularly significant since it confers resistance not only to *Leishmania* infection but also to two other unrelated pathogens viz. *Mycobacteria* and *Salmonella* (20). Although the function of Nramp1 as a phago-endosomal iron transporter is well-established, there are conflicting reports with respect to the direction of iron transport (24, 28–32). So far there has been no attempt to investigate how Nramp1 expression is regulated in an infected cell or how it controls iron availability to intracellular pathogens. Hence, the mechanism by which Nramp1 acts against the invading pathogens remains poorly understood. In this scenario, our work is the first comprehensive study of Nramp1 expression in the context of an infection. Employing macrophage infection model of *L. major,* we made an interesting observation that Nramp1 protein level undergoes time-dependent changes during the course of *Leishmania* infection. First, at 12 hours post infection, there was a drastic reduction in the level of Nramp1 which was followed by near-complete recovery of expression at 30 hours. Downregulation of Nramp1 was accompanied by increased phagolysosomal iron content and enhanced intracellular parasite growth. We also report that *Leishmania* infection-induced downregulation of Nramp1 is caused by ubiquitin-proteasomal degradation, which in turn is mediated by the iron modulatory peptide hormone hepcidin (33). Taken together, our study highlights Nramp1 as a central player in the battle for iron between the host and the pathogen where each of them tries to tinker Nramp1 protein level to modulate phagolysosomal iron content. Moreover, our study uncovers Nramp1 as a novel target of hepcidin, in addition to its well-established target ferroportin. A proposed model describing the role of this hitherto unidentified hepcidin-Nramp1 axis in regulating phagolysosomal iron level in response to *Leishmania* infection is described in Fig. 7.

**FIG 7.**
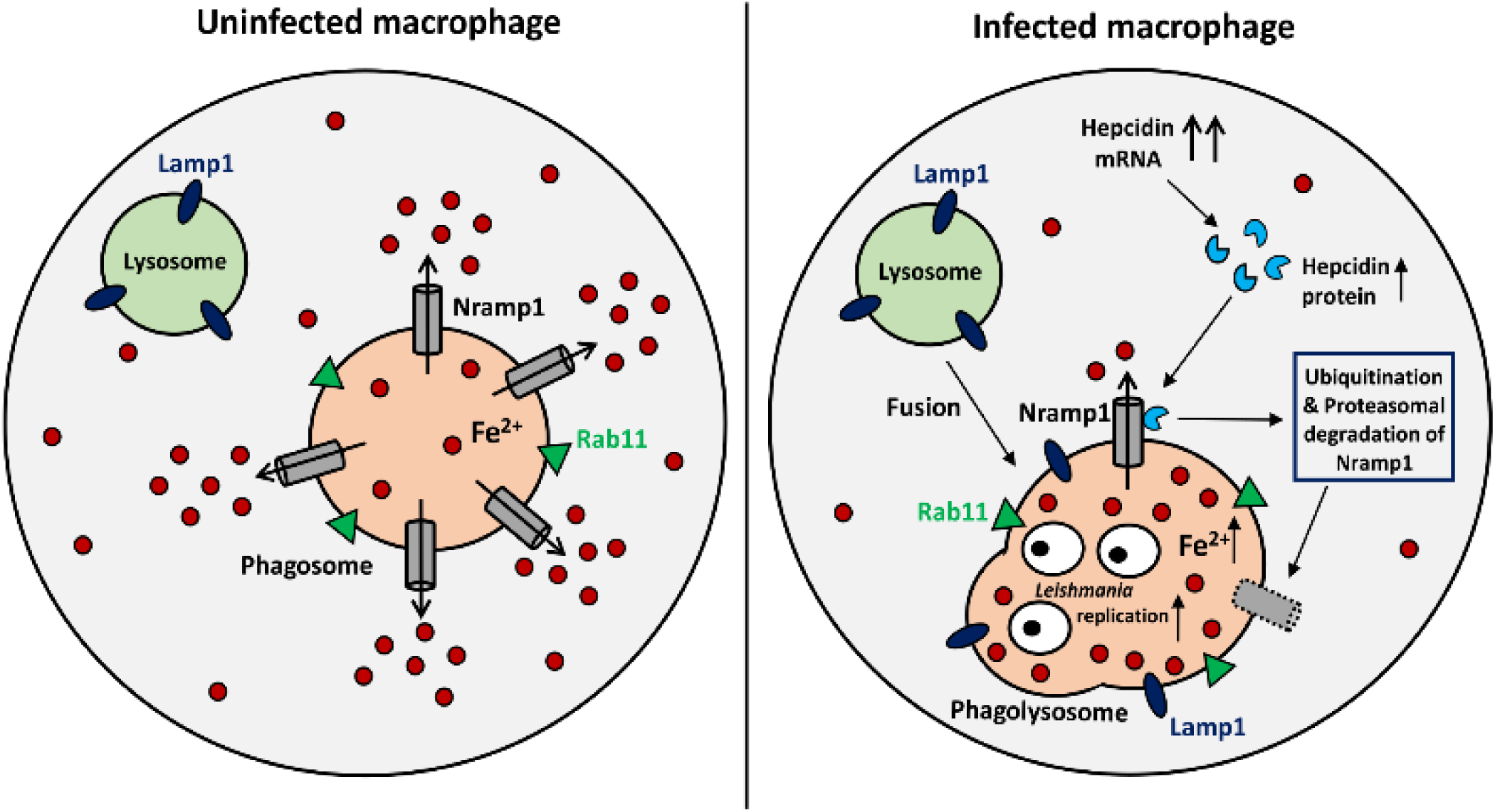
Proposed model for the *L. major* infection-induced degradation of macrophage Nramp1 and its impact on phagolysosomal iron content. Macrophage cells export iron from the phagosomal/phagolysosomal lumen to the cytosol via the action of Nramp1. While the cells become infected with *Leishmania*, it stimulates synthesis of iron regulatory peptide hormone hepcidin. In these infected cells, hepcidin binds with Nramp1, eventually causing its degradation via ubiquitine-proteasomal degradation pathway. The loss of Nramp1 results into the accumulation of iron within *Leishmania* containing phagolysosomal compartment that favors the replication of the parasite.

It was previously reported that in uninfected macrophages Nramp1 resides in the endocytic compartments, which are involved in phagolysosome biogenesis (25, 26, 45). Upon *L. major* infection, Nramp1 was found in the membrane of the pathogen-containing phagolysosomes thereby raising the question whether it can influence the parasite microenvironment by virtue of its iron transport activity (26). Corroborating evidence was provided by our immunofluorescence study that revealed recruitment of Nramp1 to the Rab11/Lamp1 double-positive phagolysosomal compartments in *L. major* infected macrophages. Unexpectedly, we also noticed a significant reduction in the Nramp1 protein expression at 12 hours post infection. This prompted us to investigate if reduced Nramp1 level in *Leishmania* infected macrophages resulted in any change in the phagolysosomal iron content. Addressing this issue was crucial for understanding the mechanism by which Nramp1 counteracts intracellular pathogens. Indeed, at 12 hours post infection, there was almost two folds increase in phagolysosomal iron level in *L. major* infected macrophages as compared to their uninfected counterparts. Our data thus established an inverse relationship between Nramp1 protein level and phagosomal/phagolysosomal iron content, which is supportive of the notion that Nramp1 transports iron from intracellular vesicles to the cytosol (29, 30). This notion was further strengthened by the observation that at 30 hours post infection Nramp1 expression reverted to the normal level with concomitant decrease in the phagosomal/phagolysosomal iron content.

Growth and survival of *Leishmania*, like other intracellular pathogens, is critically dependent on the availability of iron and their ability to scavenge it from the surroundings (34, 46, 47). Since iron pool in mammals is tightly regulated, the battle between the host and the pathogen for this essential micronutrient has often been found to be a key determinant of the infection outcome (48). Recently, Ben-Othman *et. al.* reported that *L. amazonensis* infection triggered downregulation of ferroportin, the macrophage cell surface iron exporter, as a strategy to enhance intracellular iron level and promote parasite growth (37). However, it remained unclear how this augmented cytosolic iron pool is accessed by *Leishmania*, which resides within the phagolysosomal compartment. In this regard, our work unraveled an alternative Nramp1-targetted iron-scavenging mechanism by which *Leishmania* infection increases phagolysosomal iron level that can be directly taken up by the parasite employing its own iron transporter (49–51). It was therefore not surprising that intracellular parasite count peaked at 12 hours post infection, the time at which phagolysosomal iron level was also at its maximum because of stunted Nramp1 expression. But what caused reversal of Nramp1 expression to the normal level at 30 hours post infection is somewhat mysterious. From our data it appears to be a host defense response designed to create an iron-limiting microenvironment for the invading pathogens so as to restrict their propagation.

After having established infection-induced downregulation of Nramp1 as a novel iron-sequestering strategy of *Leishmania*, attempts were made to obtain mechanistic insights of this process. Several important macrophage genes, especially those linked with immune response against pathogens, were shown to be transcriptionally repressed upon *Leishmania* infection (13–15). But the possibility of transcriptional inhibition was ruled out in this case as there was no difference in the Nramp1 transcript level between uninfected and *Leishmania* infected macrophages. Rather, Nramp1 downregulation at 12 hours post infection was found to be completely blocked by treatment with the proteasome inhibitor, MG132. This result along with the observation that Nramp1 ubiquitination was significantly enhanced in *Leishmania* infected macrophages supported the conclusion that infection-induced downregulation of Nramp1 is mediated by ubiquitin-proteasomal degradation pathway (36). There are multiple reports confirming direct involvement of *Leishmania* surface protease GP63 in degradation of host proteins as a tool to manipulate macrophage signaling and function (52–55). However, engaging the host proteasomal machinery to selectively target a host protein is somewhat unique and seems to be a clever tactic employed by the parasite to alter the phagolysosomal microenvironment. To the best of our knowledge there is only one such prior study where *L. donovani* infection was shown to subvert JAK2/STAT1α signaling pathway in macrophage through proteasomal degradation of STAT1α (56). It is worth noting that MG132-mediated stabilization of Nramp1 was accompanied with significantly reduced phagolysosomal iron content resulting in more than fifty percent lowering of intracellular parasite burden. Based on the available data it is thus tempting to speculate that inhibition of macrophage proteasome might be an effective way to target intracellular *Leishmania*. Apart from depriving the pathogen of iron, proteasome inhibition may also restore JAK2/STAT1α signaling pathway of the macrophage and activate cytokine-mediated antiparasitic immune response (57). Selective targeting of *Leishmania* proteasome has recently shown the promise to treat visceral leishmaniasis in preclinical studies (58, 59). But the possibility of mammalian proteasome-directed anti-leishmanial therapy is yet to be explored. Since an FDA approved mammalian proteasome inhibitor (bortezomib) is already available, such host-directed, multitarget therapeutic approach is worth pursuing, which may provide a new direction towards treatment of this neglected disease (60).

What triggered ubiquitination and proteasomal degradation of Nramp1 is yet to be fully understood. However, an important lead in this direction was provided by our serendipitous finding that infection-induced degradation of Nramp1 is dependent on the iron regulatory peptide hormone hepcidin. Strikingly, heparin-mediated inhibition of hepcidin transcription not only blocked Nramp1 degradation but this treatment also resulted in depletion of phagolysosomal iron pool and drastic lowering of intracellular parasite burden. In a recent report, *L. amazonensis* infection was shown to upregulate transcription of hepcidin in macrophage, which in turn caused degradation of the cell surface iron exporter ferroportin (37). Prior to this work, an extensive body of research has demonstrated that hepcidin binding triggers rapid ubiquitination of ferroportin thereby inducing its internalization and degradation (43, 44). In view of this, our co-immunoprecipitation data showing physical interaction between Nramp1 and hepcidin is quite intriguing. Whether this interaction is indeed responsible for Nramp1 ubiquitination followed by its degradation cannot be ascertained unequivocally at this point and such causal relationship needs to be validated with follow-up studies. However, this seems to be a likely possibility since: a) Nramp1 was protected from infection-induced degradation when hepcidin expression was downregulated. Therefore, hepcidin, either alone or in association with other molecular partners, must be playing a critical role in degrading Nramp1; b) Nramp1 is topologically identical to ferroportin, with 12 transmembrane domains, hence both may follow similar hepcidin-mediated degradation mechanism (40). Although hepcidin is primarily produced by the hepatocytes and acts on the macrophage ferroportin in a paracrine fashion, there are few reports confirming its endogenous expression in the macrophages in response to different infections and inflammatory stimuli (61–63). It was also reported that macrophage-produced hepcidin may act on the cell surface localized ferroportin in an autocrine fashion to sequester iron in those cells (64). But the mechanism by which hepcidin is retained within the macrophage and act on an intracellular target like Nramp1 needs to be investigated in details to better understand the functional implications of hepcidin-Nramp1 axis. Our work is an important first step towards this direction. Since Nramp1 is reported to facilitate efficient macrophage iron recycling following erythrophagocytosis, this hitherto unidentified hepcidin-Nramp1 axis may have a broader regulatory role in maintaining iron homeostasis in the phagocytic cells (65, 66).

## Materials and Methods

Unless mentioned specifically, all reagents were purchased from Sigma-Aldrich. Primers for PCR were obtained from Integrated DNA Technologies.

### Antibodies

A rabbit polyclonal antibody was raised against mouse Nramp1 using a synthetic peptide (^321^LQNYAKIFPRDN^334^) from C-terminal region of the protein as antigen (IMGENEX India custom antibody generation facility). The antibody was validated by western blot analysis using J774A.1 macrophage whole cell lysate. Anti-Lamp1 antibody (Abcam) and anti-Rab11 antibody (Santa-Cruz Biotechnology) were kind gifts of Dr. Arnab Gupta (IISER Kolkata). Rabbit polyclonal antibody raised against human hepcidin polypeptide was a generous gift of Dr. William S. Sly (Saint Louis University School of Medicine).

### Parasite and mammalian cell culture

The *L. major* strain 5ASKH was kindly provided by Dr. Subrata Adak (IICB, Kolkata). *L. major* promastigotes were cultured in M199 medium (Gibco) pH 7.2, supplemented with 15% heat-inactivated fetal bovine serum (FBS, Gibco), 23.5 mM HEPES, 0.2mM adenine, 150 µg/ml folic acid, 10 µg/ml hemin, 120 U/ml penicillin, 120 µg/ml streptomycin, and 60 µg/ml gentamicin at 26°C. J774A.1 cells (murine macrophage cell line obtained from National Center for Cell Sciences, Pune) were grown in Dulbecco’s modified Eagle’s medium (DMEM, Gibco) pH 7.4 supplemented with 2 mM L-glutamine, 100 U/ml penicillin, 100 μg/ml streptomycin, and 10% heat-inactivated FBS at 37°C in a humidified atmosphere containing 5% CO2. Cell number was quantified using a hemocytometer (10).

### Isolation of peritoneal macrophages from BALB/c mice

BALB/c mice were obtained from the National Institute of Nutrition (NIN), Hyderabad, and housed in our institutional animal facility. Experiments with these mice were conducted according to the CPCSEA guidelines and Institutional Animal Ethics Committee approved protocol. Thioglycolate elicited peritoneal macrophages were isolated from 6-8 weeks old mice as described earlier (67). Briefly, 4 days after intraperitoneal injection of 3% Brewer’s thioglycolate medium (Himedia), mice were euthanized and peritoneal macrophages were collected using 20 G needle. Thereafter the isolated macrophages were cultured in DMEM pH 7.4 supplemented with 2 mM L-glutamine, 100 U/ml penicillin, 100 μg/ml streptomycin, and 10% heat-inactivated FBS at 37°C in a humidified atmosphere containing 5% CO2. Non-adherent cells were discarded between 18-24 h. Cellular viability was determined using trypan blue dye exclusion test.

### *L. major* infection of macrophages and determination of intracellular parasite burden

Infection of J774A.1 or primary peritoneal macrophages with *L. major* promastigotes was performed as described by us previously (10). Briefly, the macrophages were activated with 100 ng/ml lipopolysaccharide (LPS) for 6 hours. *L. major* promastigotes were then added to the macrophages at a ratio of 30:1 (parasite: macrophage) and the infection was allowed to continue for 2, 6 or 12 hours, as indicated. For 30 hours infection, J774A.1 macrophages were first incubated with *L. major* promastigotes for 12 hours following which the parasites were removed and the infected macrophage cells were incubated for another 18 hours. In each case, uninfected cells similarly treated with LPS served as control. After infection, cells were washed, fixed with acetone-methanol (1:1) and mounted with anti-fade mounting medium containing DAPI (VectaShield from Vector Laboratories). Intracellular parasite burden (number of amastigotes/100 macrophages) was quantified by counting the total number of DAPI-stained nuclei of macrophages and *L. major* amastigotes in a field (at least 100 macrophages were counted from triplicate experiments). During pharmacological inhibition studies cells were either treated with 1μM MG132 (kindly provided by Dr. Partho Sarothi Ray, IISER Kolkata) or 4 µg/ml heparin.

### Immunofluorescence studies and image analysis

Macrophages grown on glass coverslips were fixed using acetone: methanol (1: 1) for 10 minutes at room temperature. After two washes with PBS, cells were permeabilized using 0.1 % triton-X 100. Cells were washed again with PBS and blocked with 0.2% gelatin for 5min at room temperature. Cells were then incubated with desired primary antibodies (anti-Nramp1 1: 50; anti-Rab11 1: 200; and anti-Lamp1 1:20) for 1.5 hours at room temperature and thereafter washed with PBS. Cells were then incubated with either of the following secondary antibodies (Molecular Probes), goat anti-rabbit Alexa fluor 488 (1: 800), goat anti-mouse Alexa fluor 568 (1: 600) or donkey anti-goat Cy3 (1: 800) and washed with PBS. Finally, the cells were mounted on anti-fade mounting medium containing DAPI and imaged in Carl Zeiss Apotome.2 microscope using 63X oil immersion objectives or in Olympus IX-81 epifluorescence microscope using either 40X or 60X objectives. In colocalization experiments Pearson’s correlation coefficient (PCC) was calculated as described previously (68). Relative fluorescence intensity was measured using microscope’s own software ZEN Blue (florescence intensities of more than 100 macrophages were measured from triplicate experiments).

### Phagosome isolation and Iron quantification

Phagosomes from both uninfected and *L. major* infected J774A.1 macrophage cells were isolated using the sucrose density gradient centrifugation as described previously (69). Isolated phagosomes were subjected to western blot to verify the presence of both Nramp1 and Rab11 in the desired fraction. The primary antibody dilutions were: anti-Nramp1 antibody (1: 500) and anti-Rab11 antibody (1: 1000). Following the overnight incubation with primary antibodies at 4°C blots were probed with HRP-conjugated goat anti-rabbit or rabbit anti-goat secondary antibodies respectively at 1: 4000 dilutions. Phagosomal iron (Fe^2+^) was quantified using ferrozine assay as reported earlier (70). Briefly, 100μL of phagosomal fraction was incubated with 100μL 10mM HCl, 4.5% KMnO4 for 2 hours at 60°C. After this samples were cooled down and further incubated with iron detection reagent (that contains 6.5mM ferrozine, 6.5mM neocuproine, 2.5M ammonium acetate and 1M ascorbic acid) for 30 mins. Thereafter, the absorbance was measured at 550nm using microplate reader. A standard curve was prepared using varying concentration of FeCl3 (0-300 μM) and the iron concentration in experimental samples was derived from the standard curve.

### Co-Immunoprecipitation assay

To assess ubiquitination status of Nramp1, whole cell lysates were subjected to immunoprecipitation (IP) with anti-Nramp1 followed by immunoblotting with anti-ubiquitin antibody. Briefly, J774A.1 macrophage cells were either uninfected or infected with *L. major* for 12 hours following which cellular proteins were extracted in lysis buffer containing 150 mM NaCl, 10mM EDTA, 10mM Tris (pH 7.4), 1% Triton X-100 and protease inhibitor cocktail. Lysates containing equal amount of protein were incubated with 60µl of Protein A PLUS Agarose Beads (BioBharati Life Science Pvt. Ltd., India) for 15min at 4°C and centrifuged at 3000rpm for 2min (71). The supernatant was incubated with 5µl of anti-Nramp1 overnight at 4°C in mild shaking. Thereafter 60µl of Protein A PLUS Agarose Bead was added to it and incubated for 4 hours at 4°C followed by centrifugation. The immunoprecipitates were washed twice with 1.0ml lysis buffer and dissolved in 100µl sample buffer. The samples were then subjected to SDF-PAGE followed by western blot using either anti-ubiquitin or Nramp1 antibody. To determine the binding of hepcidin and Nramp1 we performed immunoprecipitation using Thermo Scientific Pierce Co-IP kit (kindly provided by Dr. Piyali Mukherjee of Presidency University, Kolkata) following the manufacturer’s protocol. At first anti-Nramp1 was coupled to AminoLink Plus resin, which was then used for the immunoprecipitation assay following similar method as described above. Covalent coupling of the antibody to the resin provides an advantage over the traditional Co-IP methods that use Protein A or G resulting in co-elution of the antibody heavy and light chains. Presence of either hepcidin or Nramp1 in the immunoprecipitates was determined by immunodot blot method using the corresponding antibodies.

### Western blot and immunodot blot

Uninfected and infected J774A.1 macrophage cells were washed twice with ice cold PBS and scrapped to collect it in a centrifuge tube. Cell suspension was centrifuged at 3000 rpm for 5 min. The cell pellet was resuspended in lysis buffer containing PBS, protease inhibitor cocktail and was sonicated to prepare whole cell lysate. The whole cell lysates were subjected to SDS-PAGE and Nramp1 protein level was determined using western blot. Anti-Nramp1 antibody was used at a dilution of 1: 1000 and anti-γ-Actin antibody (Bio-Bharati) was used at a dilution of 1: 4000. HRP-conjugated goat anti-rabbit secondary antibody was used at a dilution of 1: 4000. The blots were developed using SuperSignal West Pico Chemiluminescent Substrate (Pierce). Quantification of the band intensity was performed using ImageJ software. Hepcidin levels in the whole cell lysate or immunoprecipitate were determined by immunodot blot assay as described previously (72). For this, the samples were spotted directly on the PVDF membrane and probed with anti-hepcidin, anti-Nramp1 or anti-γ-Actin primary antibodies at a dilution of 1: 2000 for each. HRP-conjugated goat anti-rabbit secondary antibody was used at a dilution of 1: 7000.

### RNA isolation and q RT-PCR

Total RNA was isolated from both uninfected and infected macrophages using TRIzol reagent (Invitrogen). These were further treated with DNaseI (Invitrogen) to remove any DNA contaminations. The cDNA was synthesized using Verso cDNA synthesis kit (Thermo) from 1µg of total RNA. Transcript level of different genes was quantified using the following primers: mouse Nramp1-Forward 5’-TTACTCACTCGGACCAGCAC-3’, mouse Nramp1 Reverse 5’-GGGGGCTCTTGTCACTAATCAT-3’; mouse Hepcidin Forward 5’-TGTCTCCTGCTTCTCCTCCT-3’, mouse hepcidin Reverse 5’-CTCTGTAGTCTGTCTCAT-3’; β-Actin Forward 5’-GGCTGTATTCCCCTCCATCG-3’, beta-Actin Reverse 5’-CCAGTTGGTAACAATGCCATG T-3’. Real time PCR was performed using 7500 real time PCR system of applied Biosystems with SYBR green fluorophore (BioRad). Transcript levels of either Nramp1 or Hepcidin were normalized with respect to β-Actin expression in each of the sample.

### Statistical analysis

All statistical analyses were executed by Student’s t test or one-way ANOVA calculator. The results were represented as mean ± SD from minimum 3 independent experiments. P values of ≤0.05 were considered statistically significant and levels of statistical significance were indicated as follows: * p≤0.05, ** p<0.01, *** p<0.001, **** p<0.0001.

## Acknowledgements

The authors sincerely thank Mr. Ritabrata Ghosh, Mr. Dipesh Dutta and Mr. Sujoy Bose for their expert technical assistance. Drs. William S. Sly, Abdul Waheed and Piyali Mukherjee are acknowledged for helpful suggestions and for providing critical reagents used in this work.

This work was supported by IISER Kolkata intramural fund. S. Banerjee was supported by DST INSPIRE PhD fellowship.

The authors declare no competing financial interests.

## Author contributions

S. Banerjee designed and performed the experiments, analyzed the data and assisted in manuscript writing. R. Datta conceptualized and supervised the work, analyzed the data, acquired funds and wrote the manuscript.

## Abbreviations

Nramp1: Natural resistance associated macrophage protein 1
Slc11a1: Solute carrier family 11 member 1
*L. major*: *Leishmania major*

## Figure Legends

**FIG S1.**
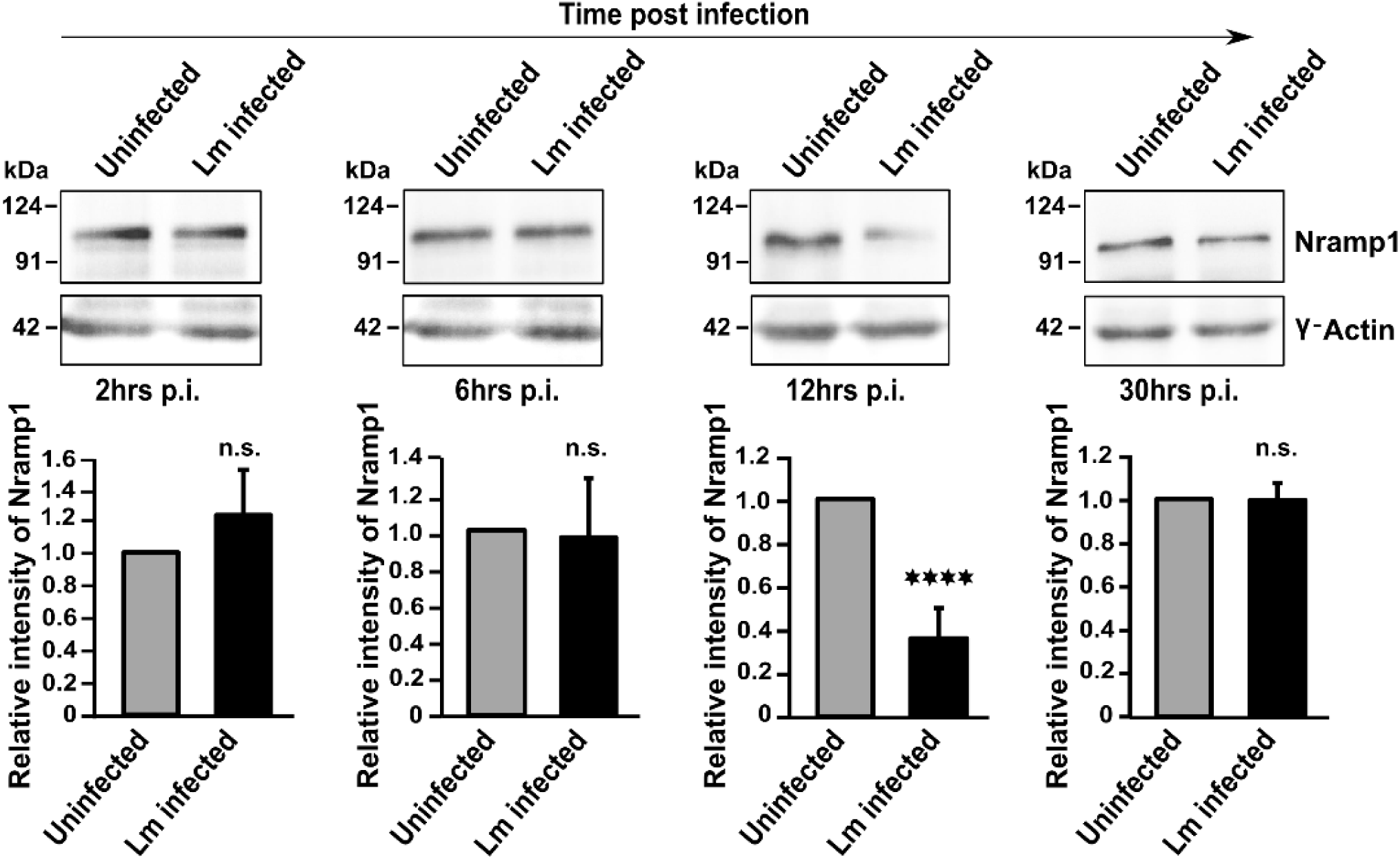
*L. major* infection-induced downregulation of Nramp1 at 12hrs post infection. J774A.1 macrophage cells were either uninfected or infected with *L. major* (Lm) for 2-30 hours. After indicated time points post infection (p.i.), cells were lysed and Nramp1 protein level was determined in whole cell lysates by western blot. Blots were probed with anti-Nramp1 and anti-γ-Actin (loading control) antibody where both Namp1 and γ-Actin bands appeared at their predicted molecular weight of ∼100 and 42 kDa respectively. Lower panel shows the relative intensity of Nramp1 protein in uninfected (grey bar) and Lm infected (black bar) macrophages measured using ImageJ software. During quantification uninfected cell was used as reference sample and γ-Actin as endogenous control protein. Error bars represent SEM values calculated from at least three independent experiments and normalized to respective control. ****, p≤0.0001.

**FIG S2.**
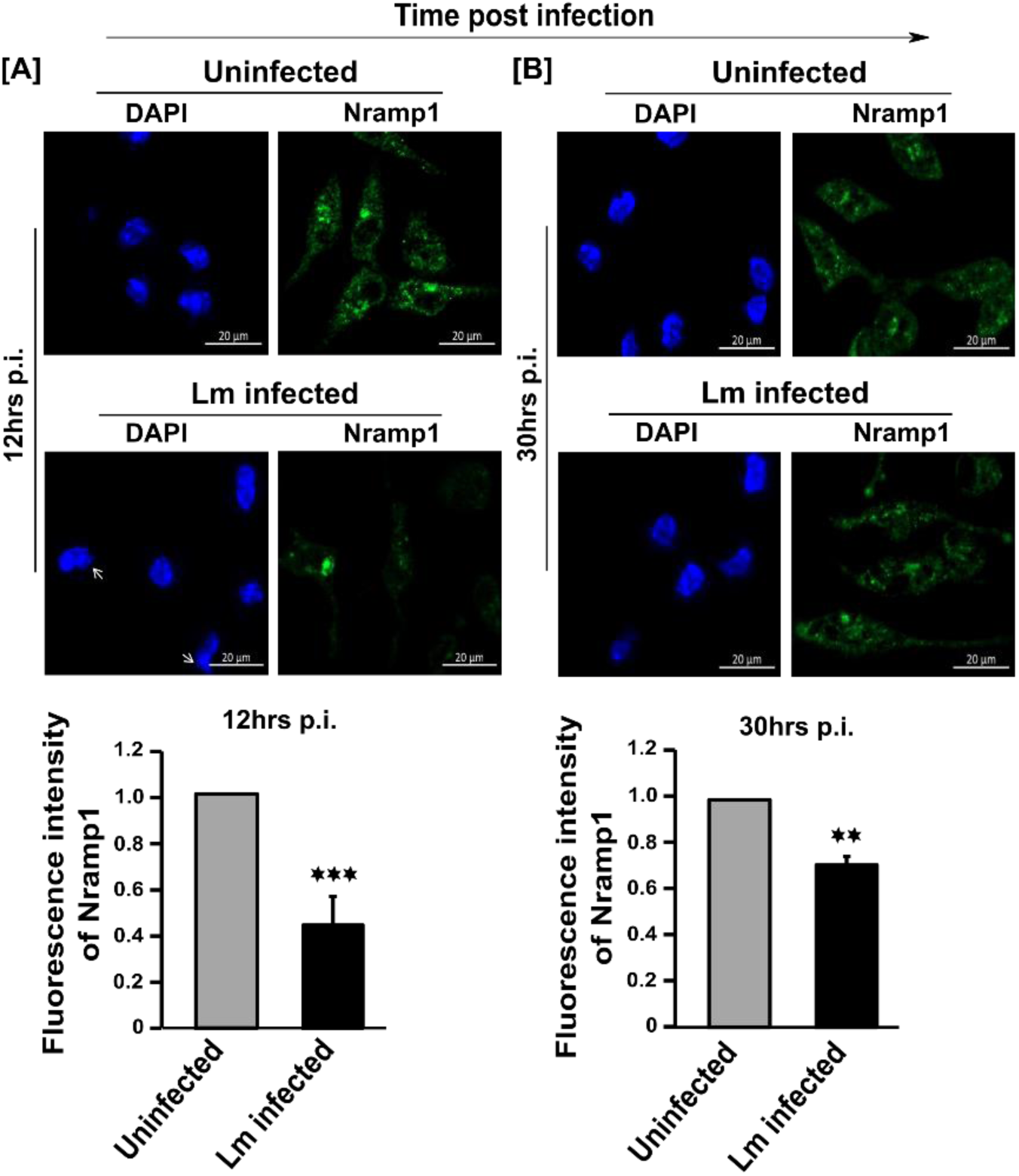
Nramp1 protein level is reduced in *L. major* infected peritoneal macrophages. (A and B) BALB/c mice derived thioglycollate elicited peritoneal macrophages were either uninfected or infected with *L. major* (Lm) promastigotes for 12hrs (A) or 30hrs (B) and Nramp1 protein level was examined by immunostaining using anti-Nramp1 antibody (Green). *Leishmania* and macrophage nucleus was stained with DAPI (Blue). Arrows indicate the presence of intracellular parasites in Lm infected macrophages. Cells were visualized with Zeiss Apotome microscope using 63X oil immersion objective. Scale bar: 20µm. Lower panel shows the relative fluorescence intensity of Nramp1 in uninfected (grey bar) and Lm infected (black bar) macrophages at both the time points. Relative fluorescence intensity was measured from at least 100 macrophage cells imaged under Olympus IX-81 epifluorescence microscope from three independent experiments using ZEN software. Uninfected cells were used as reference sample. Error bars represent SEM values calculated from the experiments. **, p≤0.01; ***, p≤0.001.

**FIG S3.**
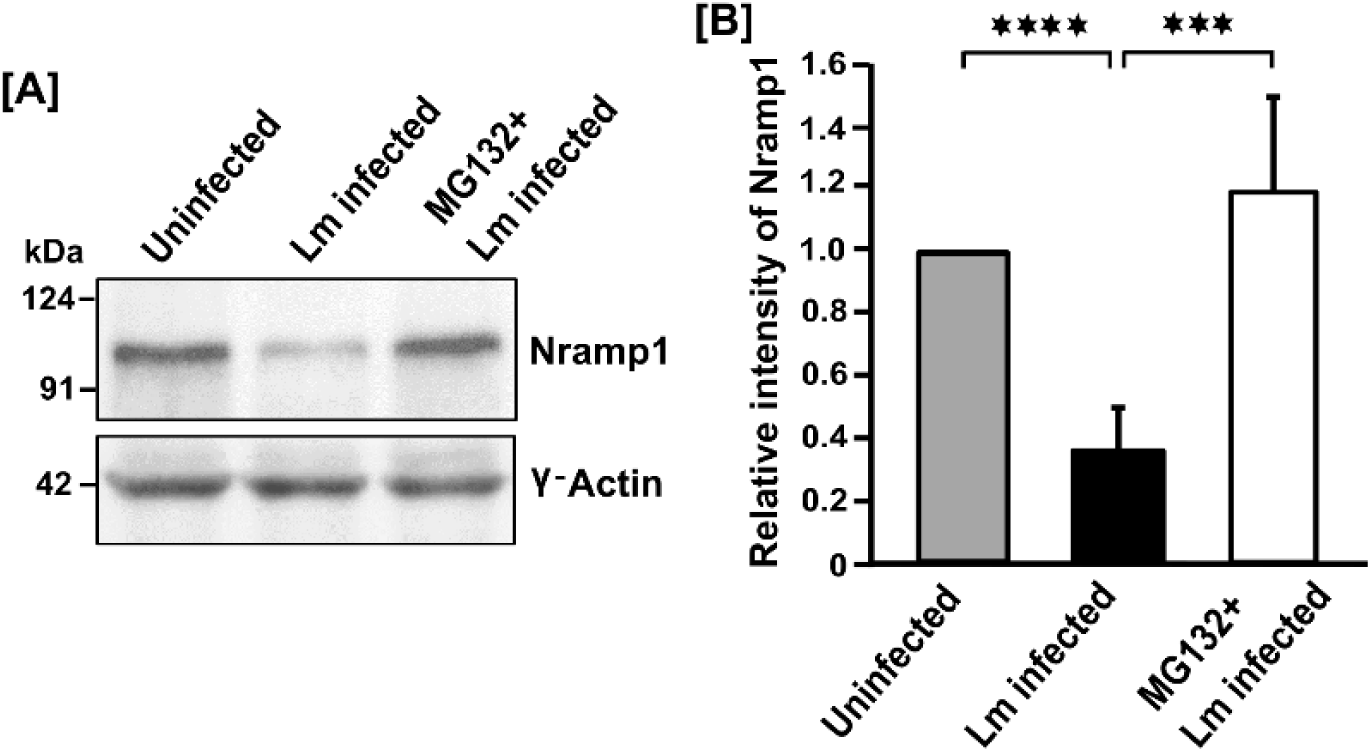
Pharmacological inhibition of proteasome activity prevents *L. major* infection-induced degradation of Nramp1. (A) Uninfected, *L. major* (Lm) infected or 1µM MG132 treated Lm infected J774A.1 macrophage cells were lysed at 12hrs post infection and Nramp1 protein level was determined by western blot. Blots were probed with anti-Nramp1 and anti-γ-Actin (as loading control) antibodies. (B) Bar diagram showing relative intensity of Nramp1 protein in uninfected (grey), Lm infected (black) and 1µM MG132 treated Lm infected (white) J774A.1 macrophages measured from western blot data using ImageJ software as mentioned earlier. Error bars represent SEM values calculated from at least three independent experiments. ***, p≤0.001; ****, p≤0.0001.

**FIG S4.**
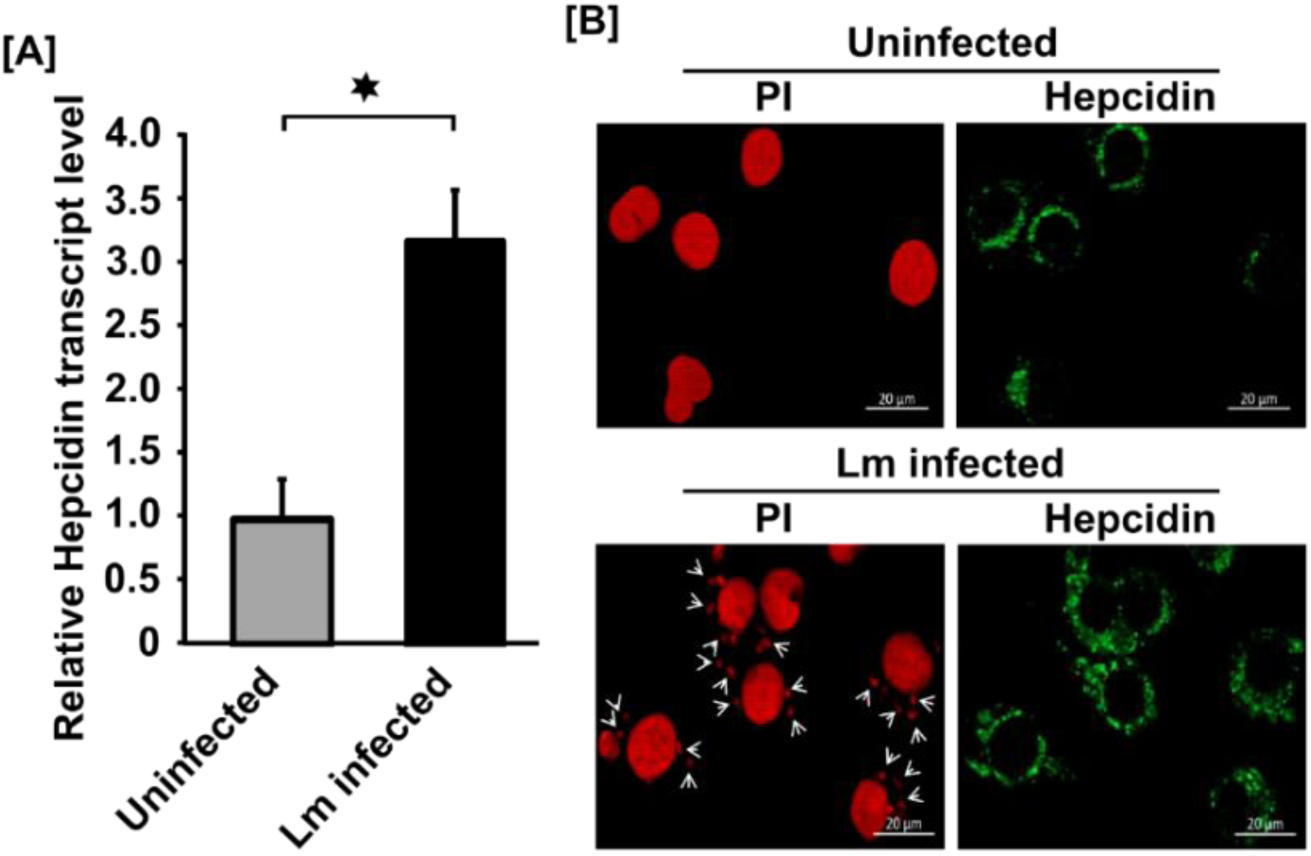
*L. major* infection stimulates hepcidin expression in macrophage both at mRNA at protein level. (A) J774A.1 macrophage cells were either uninfected or infected with *L. major* (Lm) promastigotes for 12hrs and cDNA was been prepared from each of the samples. The hepcidin mRNA level was then measured by RT-qPCR using β-Actin as endogenous control and uninfected cells as reference sample. Error bars represent SEM values calculated from three independent experiments. *, p≤0.05. (B) Uninfected or Lm infected J774A.1 macrophage cells at 12hrs post infection were fixed and immunostained using anti-hepcidin antibody (green). Propidium iodide (PI, red) was used to stain parasite and host cell nucleus. Arrows indicate the presence of intracellular parasites in Lm infected macrophages. Cells were visualized with Zeiss Apotome microscope using 63X oil immersion objective. Scale bar: 20µm.

**FIG S5.**
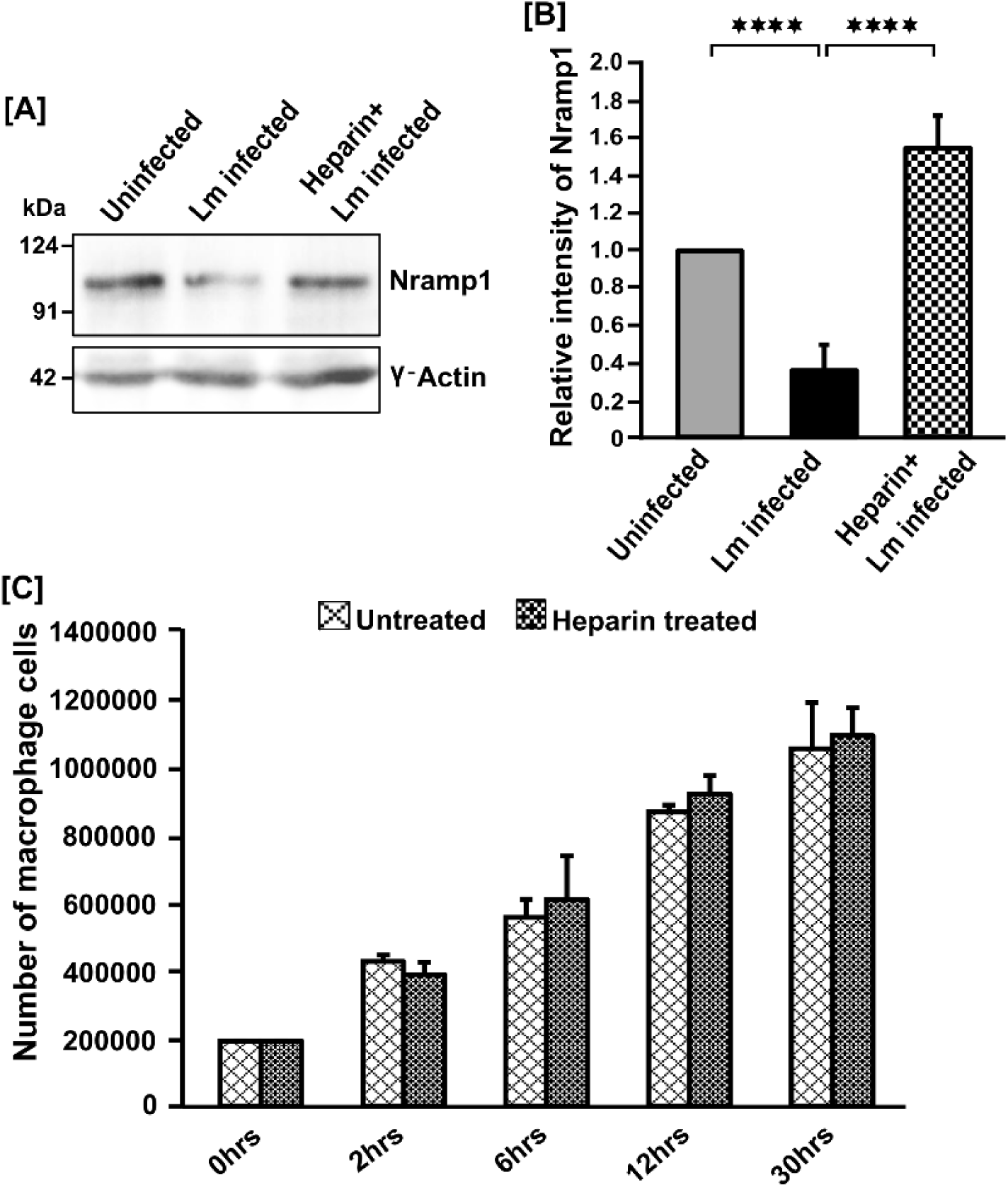
Pharmacological inhibition of hepcidin transcription prevents Nramp1 degradation without effecting cell viability. (A) Uninfected, *L. major* (Lm) infected or 4µg/ml heparin treated Lm infected J774A.1 macrophage cells were lysed at 12hrs post infection and Nramp1 protein level was analyzed by western blot using anti-Nramp1 and γ-Actin antibody (as loading control). (B) Bar diagram shows relative intensity of Nramp1 protein in uninfected (grey), Lm infected (black) and 4µg/ml heparin treated Lm infected (checker board pattern) J774A.1 macrophages measured from the respective western blot data using ImageJ software. All the measurements were normalized using uninfected cells as reference sample and γ-Actin as endogenous control. Error bars represent SEM values calculated from three independent experiments. ***, p≤0.001; ****, p≤0.0001. (C) J774A.1 macrophage cells were grown in presence of 4µg/ml heparin for 0-30hrs and cellular viability was measured by trypan blue dye exclusion test. Cells were observed under 40X magnification of inverted light microscope (Nikon). Error bars represent SEM values calculated from three independent experiments.

## References

1. Burza S, Croft SL, Boelaert M. 2018. Leishmaniasis. Lancet (London, England) 392:951–970.

2. Okwor I, Uzonna J. 2016. Social and Economic Burden of Human Leishmaniasis. Am J Trop Med Hyg 94:489–493.

3. Croft SL, Sundar S, Fairlamb AH. 2006. Drug resistance in leishmaniasis. Clin Microbiol Rev 19:111–26.

4. Peters NC, Egen JG, Secundino N, Debrabant A, Kimblin N, Kamhawi S, Lawyer P, Fay MP, Germain RN, Sacks D. 2008. In vivo imaging reveals an essential role for neutrophils in leishmaniasis transmitted by sand flies. Science 321:970–4.

5. Chang KP, Dwyer DM. 1978. Leishmania donovani. Hamster macrophage interactions in vitro: cell entry, intracellular survival, and multiplication of amastigotes. J Exp Med 147:515–30.

6. Thi EP, Lambertz U, Reiner NE. 2012. Sleeping with the Enemy: How Intracellular Pathogens Cope with a Macrophage Lifestyle. PLoS Pathog 8:e1002551.

7. Haas A. 2007. The Phagosome: Compartment with a License to Kill. Traffic 8:311–330.

8. Burchmore RJ, Barrett MP. 2001. Life in vacuoles--nutrient acquisition by Leishmania amastigotes. Int J Parasitol 31:1311–20.

9. Van Assche T, Deschacht M, da Luz RAI, Maes L, Cos P. 2011. Leishmania–macrophage interactions: Insights into the redox biology. Free Radic Biol Med 51:337–351.

10. Pal DS, Abbasi M, Mondal DK, Varghese BA, Paul R, Singh S, Datta R. 2017. Interplay between a cytosolic and a cell surface carbonic anhydrase in pH homeostasis and acid tolerance of Leishmania. J Cell Sci 130:754–766.

11. Späth GF, Garraway LA, Turco SJ, Beverley SM. 2003. The role(s) of lipophosphoglycan (LPG) in the establishment of Leishmania major infections in mammalian hosts. Proc Natl Acad Sci U S A 100:9536–41.

12. Gregory DJ, Sladek R, Olivier M, Matlashewski G. 2008. Comparison of the Effects of Leishmania major or Leishmania donovani Infection on Macrophage Gene Expression. Infect Immun 76:1186–1192.

13. Beverley SM. 1996. Hijacking the cell: parasites in the driver’s seat. Cell 87:787–9.

14. Carrera L, Gazzinelli RT, Badolato R, Hieny S, Muller W, Kuhn R, Sacks DL. 1996. Leishmania promastigotes selectively inhibit interleukin 12 induction in bone marrow-derived macrophages from susceptible and resistant mice. J Exp Med 183:515–526.

15. Buates S, Matlashewski G. 2001. General Suppression of Macrophage Gene Expression During *Leishmania donovani* Infection. J Immunol 166:3416–3422.

16. Sakthianandeswaren A, Foote SJ, Handman E. 2009. The role of host genetics in leishmaniasis. Trends Parasitol 25:383–391.

17. Jeronimo SMB, Duggal P, Ettinger NA, Nascimento ET, Monteiro GR, Cabral AP, Pontes NN, Lacerda HG, Queiroz PV, Gomes CEM, Pearson RD, Blackwell JM, Beaty TH, Wilson ME. 2007. Genetic Predisposition to Self-Curing Infection with the Protozoan *Leishmania chagasi:* A Genomewide Scan. J Infect Dis 196:1261–1269.

18. Blackwell JM, Fakiola M, Ibrahim ME, Jamieson SE, Jeronimo SB, Miller EN, Mishra A, Mohamed HS, Peacock CS, Raju M, Sundar S, Wilson ME. 2009. Genetics and visceral leishmaniasis: of mice and man. Parasite Immunol 31:254–66.

19. Mohamed HS, Ibrahim ME, Miller EN, Peacock CS, Khalil EAG, Cordell HJ, Howson JMM, El Hassan AM, Bereir REH, Blackwell JM. 2003. Genetic susceptibility to visceral leishmaniasis in The Sudan: linkage and association with IL4 and IFNGR1. Genes Immun 4:351–355.

20. Vidal SM, Malo D, Vogan K, Skamene E, Gros P. 1993. Natural resistance to infection with intracellular parasites: isolation of a candidate for Bcg. Cell 73:469–85.

21. Bucheton B, Abel L, Kheir MM, Mirgani A, El-Safi SH, Chevillard C, Dessein A. 2003. Genetic control of visceral leishmaniasis in a Sudanese population: candidate gene testing indicates a linkage to the NRAMP1 region. Genes Immun 4:104–109.

22. Mohamed HS, Ibrahim ME, Miller EN, White JK, Cordell HJ, Howson JMM, Peacock CS, Khalil EAG, El Hassan AM, Blackwell JM. 2004. SLC11A1 (formerly NRAMP1) and susceptibility to visceral leishmaniasis in The Sudan. Eur J Hum Genet 12:66–74.

23. Fattahi-Dolatabadi M, Mousavi T, Mohammadi-Barzelighi H, Irian S, Bakhshi B, Nilforoushzadeh M-A, Shirani-Bidabadi L, Hariri M-M, Ansari N, Akbari N. 2016. NRAMP1 gene polymorphisms and cutaneous leishmaniasis: An evaluation on host susceptibility and treatment outcome. J Vector Borne Dis 53:257–63.

24. Gruenheid S, Skamene E, Gros P. 1999. Nramp1: A novel macrophage protein with a key function in resistance to intracellular pathogens. Adv Cell Mol Biol Membr Organelles 5:345–362.

25. Gruenheid S, Pinner E, Desjardins M, Gros P. 1997. Natural resistance to infection with intracellular pathogens: the Nramp1 protein is recruited to the membrane of the phagosome. J Exp Med 185:717–30.

26. Searle S, Bright NA, Roach TI, Atkinson PG, Barton CH, Meloen RH, Blackwell JM. 1998. Localisation of Nramp1 in macrophages: modulation with activation and infection. J Cell Sci 111 (Pt 19):2855–66.

27. Vidal SM, Pinner E, Lepage P, Gauthier S, Gros P. 1996. Natural resistance to intracellular infections: Nramp1 encodes a membrane phosphoglycoprotein absent in macrophages from susceptible (Nramp1 D169) mouse strains. J Immunol 157:3559–68.

28. Atkinson PG, Barton CH. 1998. Ectopic expression of Nramp1 in COS-1 cells modulates iron accumulation. FEBS Lett 425:239–42.

29. Atkinson PG, Barton CH. 1999. High level expression of Nramp1G169 in RAW264.7 cell transfectants: analysis of intracellular iron transport. Immunology 96:656–62.

30. Barton CH, Biggs TE, Baker ST, Bowen H, Atkinson PGP. 1999. *Nramp1* : a link between intracellular iron transport and innate resistance to intracellular pathogens. J Leukoc Biol 66:757–762.

31. Kuhn DE, Baker BD, Lafuse WP, Zwilling BS. 1999. Differential iron transport into phagosomes isolated from the RAW264.7 macrophage cell lines transfected with Nramp1^Gly169^ or Nramp1^Asp169^. J Leukoc Biol 66:113–119.

32. Zwilling BS, Kuhn DE, Wikoff L, Brown D, Lafuse W. 1999. Role of iron in Nramp1-mediated inhibition of mycobacterial growth. Infect Immun 67:1386–92.

33. Drakesmith H, Prentice AM. 2012. Hepcidin and the Iron-Infection Axis. Science (80-) 338:768–772.

34. Flannery AR, Renberg RL, Andrews NW. 2013. Pathways of iron acquisition and utilization in Leishmania. Curr Opin Microbiol 16:716–721.

35. Lee DH, Goldberg AL. 1998. Proteasome inhibitors: valuable new tools for cell biologists. Trends Cell Biol 8:397–403.

36. Collins GA, Goldberg AL. 2017. The Logic of the 26S Proteasome. Cell 169:792–806.

37. Ben-Othman R, Flannery AR, Miguel DC, Ward DM, Kaplan J, Andrews NW. 2014. Leishmania-Mediated Inhibition of Iron Export Promotes Parasite Replication in Macrophages. PLoS Pathog 10:e1003901.

38. Chang K, Dwyer D. 1976. Multiplication of a human parasite (Leishmania donovani) in phagolysosomes of hamster macrophages in vitro. Science (80-) 193:678–680.

39. McConville MJ, de Souza D, Saunders E, Likic VA, Naderer T. 2007. Living in a phagolysosome; metabolism of Leishmania amastigotes. Trends Parasitol 23:368–375.

40. Wallace DF, Harris JM, Subramaniam VN. 2010. Functional analysis and theoretical modeling of ferroportin reveals clustering of mutations according to phenotype. Am J Physiol Physiol 298:C75–C84.

41. Nemeth E, Ganz T. 2006. Regulation of Iron Metabolism by Hepcidin. Annu Rev Nutr 26:323–342.

42. Poli M, Girelli D, Campostrini N, Maccarinelli F, Finazzi D, Luscieti S, Nai A, Arosio P. 2011. Heparin: a potent inhibitor of hepcidin expression in vitro and in vivo. Blood 117:997–1004.

43. Nemeth E, Tuttle MS, Powelson J, Vaughn MB, Donovan A, Ward DM, Ganz T, Kaplan J. 2004. Hepcidin Regulates Cellular Iron Efflux by Binding to Ferroportin and Inducing Its Internalization. Science (80-) 306:2090–2093.

44. Qiao B, Sugianto P, Fung E, del-Castillo-Rueda A, Moran-Jimenez M-J, Ganz T, Nemeth E. 2012. Hepcidin-Induced Endocytosis of Ferroportin Is Dependent on Ferroportin Ubiquitination. Cell Metab 15:918–924.

45. Desjardins M, Huber LA, Parton RG, Griffiths G. 1994. Biogenesis of phagolysosomes proceeds through a sequential series of interactions with the endocytic apparatus. J Cell Biol 124:677–688.

46. Mittra B, Cortez M, Haydock A, Ramasamy G, Myler PJ, Andrews NW. 2013. Iron uptake controls the generation of Leishmania infective forms through regulation of ROS levels. J Exp Med 210:401–16.

47. Das NK, Biswas S, Solanki S, Mukhopadhyay CK. 2009. *Leishmania donovani* depletes labile iron pool to exploit iron uptake capacity of macrophage for its intracellular growth. Cell Microbiol 11:83–94.

48. Sutak R, Lesuisse E, Tachezy J, Richardson DR. 2008. Crusade for iron: iron uptake in unicellular eukaryotes and its significance for virulence. Trends Microbiol 16:261–8.

49. Jacques I, Andrews NW, Huynh C. 2010. Functional characterization of LIT1, the Leishmania amazonensis ferrous iron transporter. Mol Biochem Parasitol 170:28–36.

50. Flannery AR, Huynh C, Mittra B, Mortara RA, Andrews NW. 2011. LFR1 Ferric Iron Reductase of *Leishmania amazonensis* Is Essential for the Generation of Infective Parasite Forms. J Biol Chem 286:23266–23279.

51. Huynh C, Sacks DL, Andrews NW. 2006. A Leishmania amazonensis ZIP family iron transporter is essential for parasite replication within macrophage phagolysosomes. J Exp Med 203:2363–2375.

52. Gomez MA, Contreras I, Halle M, Tremblay ML, McMaster RW, Olivier M. 2009. Leishmania GP63 Alters Host Signaling Through Cleavage-Activated Protein Tyrosine Phosphatases. Sci Signal 2:ra58–ra58.

53. Contreras I, Gómez MA, Nguyen O, Shio MT, McMaster RW, Olivier M. 2010. Leishmania-Induced Inactivation of the Macrophage Transcription Factor AP-1 Is Mediated by the Parasite Metalloprotease GP63. PLoS Pathog 6:e1001148.

54. Hallé M, Gomez MA, Stuible M, Shimizu H, McMaster WR, Olivier M, Tremblay ML. 2009. The *Leishmania* Surface Protease GP63 Cleaves Multiple Intracellular Proteins and Actively Participates in p38 Mitogen-activated Protein Kinase Inactivation. J Biol Chem 284:6893–6908.

55. Isnard A, Shio MT, Olivier M. 2012. Impact of Leishmania metalloprotease GP63 on macrophage signaling. Front Cell Infect Microbiol 2:72.

56. Forget G, Gregory DJ, Olivier M. 2005. Proteasome-mediated degradation of STAT1alpha following infection of macrophages with Leishmania donovani. J Biol Chem 280:30542–9.

57. Murray PJ. 2007. The JAK-STAT signaling pathway: input and output integration. J Immunol 178:2623–9.

58. Wyllie S, Brand S, Thomas M, De Rycker M, Chung C-W, Pena I, Bingham RP, Bueren-Calabuig JA, Cantizani J, Cebrian D, Craggs PD, Ferguson L, Goswami P, Hobrath J, Howe J, Jeacock L, Ko E-J, Korczynska J, MacLean L, Manthri S, Martinez MS, Mata-Cantero L, Moniz S, Nühs A, Osuna-Cabello M, Pinto E, Riley J, Robinson S, Rowland P, Simeons FRC, Shishikura Y, Spinks D, Stojanovski L, Thomas J, Thompson S, Viayna Gaza E, Wall RJ, Zuccotto F, Horn D, Ferguson MAJ, Fairlamb AH, Fiandor JM, Martin J, Gray DW, Miles TJ, Gilbert IH, Read KD, Marco M, Wyatt PG. 2019. Preclinical candidate for the treatment of visceral leishmaniasis that acts through proteasome inhibition. Proc Natl Acad Sci U S A 116:9318–9323.

59. Khare S, Nagle AS, Biggart A, Lai YH, Liang F, Davis LC, Barnes SW, Mathison CJN, Myburgh E, Gao M-Y, Gillespie JR, Liu X, Tan JL, Stinson M, Rivera IC, Ballard J, Yeh V, Groessl T, Federe G, Koh HXY, Venable JD, Bursulaya B, Shapiro M, Mishra PK, Spraggon G, Brock A, Mottram JC, Buckner FS, Rao SPS, Wen BG, Walker JR, Tuntland T, Molteni V, Glynne RJ, Supek F. 2016. Proteasome inhibition for treatment of leishmaniasis, Chagas disease and sleeping sickness. Nature 537:229–233.

60. Kane RC, Bross PF, Farrell AT, Pazdur R. 2003. Velcade: U.S. FDA approval for the treatment of multiple myeloma progressing on prior therapy. Oncologist 8:508–13.

61. Peyssonnaux C, Zinkernagel AS, Datta V, Lauth X, Johnson RS, Nizet V. 2006. TLR4-dependent hepcidin expression by myeloid cells in response to bacterial pathogens. Blood 107:3727.

62. Nguyen N-B, Callaghan KD, Ghio AJ, Haile DJ, Yang F. 2006. Hepcidin expression and iron transport in alveolar macrophages. Am J Physiol Cell Mol Physiol 291:L417–L425.

63. Sow FB, Florence WC, Satoskar AR, Schlesinger LS, Zwilling BS, Lafuse WP. 2007. Expression and localization of hepcidin in macrophages: a role in host defense against tuberculosis. J Leukoc Biol 82:934–945.

64. Theurl I, Theurl M, Seifert M, Mair S, Nairz M, Rumpold H, Zoller H, Bellmann-Weiler R, Niederegger H, Talasz H, Weiss G. 2008. Autocrine formation of hepcidin induces iron retention in human monocytes. Blood 111:2392–2399.

65. Soe-Lin S, Apte SS, Andriopoulos B, Andrews MC, Schranzhofer M, Kahawita T, Garcia-Santos D, Ponka P. 2009. Nramp1 promotes efficient macrophage recycling of iron following erythrophagocytosis in vivo. Proc Natl Acad Sci 106:5960–5965.

66. Soe-Lin S, Sheftel AD, Wasyluk B, Ponka P. 2008. Nramp1 equips macrophages for efficient iron recycling. Exp Hematol 36:929–937.

67. Zhang X, Goncalves R, Mosser DM. 2008. The Isolation and Characterization of Murine Macrophages, p. 14.1.1–14.1.14. *In* Current Protocols in Immunology. John Wiley & Sons, Inc., Hoboken, NJ, USA.

68. Mcdonald JH, Dunn KW. 2013. Statistical tests for measures of colocalization in biological microscopy. J Microsc 252:295–302.

69. Mukherjee K, Parashuraman S, Krishnamurthy G, Majumdar J, Yadav A, Kumar R, Basu SK, Mukhopadhyay A. 2002. Diverting intracellular trafficking of Salmonella to the lysosome through activation of the late endocytic Rab7 by intracellular delivery of muramyl dipeptide. J Cell Sci 115:3693–701.

70. Riemer J, Hoepken HH, Czerwinska H, Robinson SR, Dringen R. 2004. Colorimetric ferrozine-based assay for the quantitation of iron in cultured cells. Anal Biochem 331:370–375.

71. Lowry OH, Rosebrough NJ, Farr AL, Randall RJ. 1951. Protein measurement with the Folin phenol reagent. J Biol Chem 193:265–75.

72. Schwarz P, Strnad P, von Figura G, Janetzko A, Krayenbühl P, Adler G, Kulaksiz H. 2011. A novel monoclonal antibody immunoassay for the detection of human serum hepcidin. J Gastroenterol 46:648–656.

